# Witnessing others’ suffering is associated with divergent mitochondrial transcriptional programs in human monocytes

**DOI:** 10.64898/2026.11.23.762555

**Authors:** Byeol Kim Lux, Melanie C. Kos, Xin Li, Fred W. Kolling, Martin Picard, Claudia V. Jakubzick, Tor D. Wager

## Abstract

Witnessing others’ suffering can elicit a physiological stress response, yet its effects on the broader transcriptional programs of human immune cells remain poorly characterized. To characterize these responses at the cellular level, we examined monocyte transcriptional responses to witnessing others’ suffering in 55 participants using a counterbalanced, within-subject experiment. Participants viewed documentaries depicting animal suffering (observed trauma, OT) or positive human-animal interactions (observed community, OC) on separate days, with circulating monocytes collected before and after each viewing for RNA sequencing. Mitochondrial DNA-encoded protein-coding transcripts (mt-mRNAs) showed greater pre-to-post increases following OT than OC, whereas nine nuclear-encoded mitochondrial pathways (MitoPathways), involved in metabolism, detoxification, and signaling, showed lower post-viewing expression under OT. The association between baseline and post-viewing expression was also weaker under OT for the mt-mRNAs and selected MitoPathways, suggesting baseline-dependent transcriptional responses. Genome-wide principal component analysis identified a stable axis of transcriptional variation positively associated with mt-mRNA changes and negatively associated with the nuclear-encoded MitoPathways that showed the opposite pattern with mt-mRNAs. Genes at opposite ends of this axis were enriched for intracellular molecular turnover and metabolic regulation versus cellular trafficking and signaling, extending beyond canonical adversity-related inflammatory signatures. Together, these findings reveal divergent mtDNA- and nuclear-encoded mitochondrial transcriptional patterns embedded within broader variation in monocyte gene expression, linking the experience of witnessing others’ suffering to coordinated changes in innate immune cell transcription.

## Introduction

Social species are sensitive to the distress of others, a capacity that supports threat detection, caregiving, and prosocial behavior (*1*). This sensitivity extends beyond behavioral or psychological responses. Social interaction can influence metabolic regulation and the efficiency of physiological homeostasis (*2*), while witnessing others’ distress can engage central, autonomic, neuroendocrine, and immune systems (*3–5*). Adverse social conditions, including loneliness, caregiving, bereavement, and discrimination, have also been associated with changes in leukocyte gene expression (*6*). Animal studies further suggest that witnessing social defeat can induce behavior changes and engage peripheral immune responses, including alterations in circulating inflammatory cytokines (*7*, *8*). However, how witnessing others’ suffering is associated with acute cellular responses in the human peripheral immune system remains poorly understood.

Monocytes are innate immune cells involved in response to injury and infection and are responsive to neural and endocrine signals through which psychological states can influence the peripheral immunity. These cells are continuously replenished from bone marrow, express receptors for sympathetic and neuroendocrine mediators, and can reorganize transcription within hours in response to social stimuli (*9*). Monocytes also exhibit substantial metabolic activity and bioenergetic reserve capacity (*10*). Physiological adaptation involves the allocation of metabolic resources across competing cellular demands (*11*). Mitochondria contribute to these processes through energy metabolism and cellular stress signaling. Together, these characteristics make mitochondrial gene regulation a relevant dimension of monocyte responses to physiological and psychological challenges.

Mitochondria processes depend on the coordinated contributions of two genomes: mitochondrial DNA (mtDNA) and nuclear DNA. The human mitochondrial genome includes 13 protein-coding genes involved in oxidative phosphorylation, whereas the vast majority of mitochondrial proteins are encoded by the nuclear genome. Although mitochondrial functions depend on gene products from both genomes, their transcription is governed by distinct regulatory mechanisms and need not exhibit identical patterns of change. Studies of human tissues have revealed tissue-dependent relationships between mtDNA-encoded and nuclear-encoded respiratory gene expression (*12*), while experimental mitochondrial perturbations can engage coordinated mitonuclear stress responses (*13*, *14*). These findings suggest that relationships between mitochondrial and nuclear transcription may provide insight into cellular adaptation. However, whether acute psychosocial experiences are associated with coordinated or divergent transcriptional responses across these two genomes in human immune cells remains unclear.

Transcriptomic profiling provides a cell-specific molecular readout of changes in cellular programs associated with environmental experiences. Unlike the relatively stable genome, gene-expression profiles can change over hours and differ across cell types, reflecting ongoing biological responses to environmental conditions. Social genomics research has established links between psychosocial adversity and leukocyte transcription, particularly through the conserved transcriptional response to adversity (CTRA), characterized by increased inflammatory and reduced antiviral and antibody-related gene expression (*6*, *15*). Subsequent work has implicated monocytes in these transcriptional patterns (*16*, *17*). Much of this work has focused on chronic or persistent adversity, often comparing individuals with different exposure histories. Examining acute transcriptional responses to distinct stressors within individuals could help identify underlying biological mechanisms through which repeated adversity or stress influence physiology over time. Combining targeted analyses of established gene signatures with genome-wide transcriptomic approaches may further reveal broader cellular programs through which psychosocial experiences are associated with physiological regulation.

Witnessing others’ suffering offers a unique context for examining vicarious physiological responses to socially and evolutionarily relevant information, in the absence of direct threat to the observer (*18*, *19*). Such experiences may elicit not only negative affect but also empathic concern and moral appraisal, particularly when the suffering involves vulnerable others (*20*). Experimental evidence suggests that socially meaningful emotions, like shame, can influence peripheral immune activity (*21*, *22*). Documentary depictions of animal suffering provide a naturalistic context in which these emotional and morally salient dimensions unfold within a sustained narrative. Unlike studies focusing primarily on chronic adversity or direct stress exposure, this paradigm offers an opportunity to examine acute peripheral immune responses to witnessing suffering. Yet, whether such experiences are accompanied by acute changes in peripheral immune cell gene expression remains poorly understood.

Here, we investigated acute monocyte transcriptional responses to witnessing animal suffering using a counterbalanced within-subject design. Participants watched documentary films depicting either animal suffering (observed trauma; OT) or positive human-animal interactions (observed community; OC), with circulating monocytes enriched from whole blood collected before and after each viewing session for RNA sequencing. Building on our previous characterization of emotional, autonomic, neural, and targeted cytokine-related transcriptional responses to this paradigm (*20*), we examined established CTRA gene signatures, mtDNA-encoded transcripts, and nuclear-encoded mitochondrial pathways. We further applied genome-wide principal component analysis to characterize coordinated transcriptional variation across individuals. We found partial correspondence with established CTRA signatures, alongside greater pre-to-post increases in mtDNA-encoded transcripts under OT than OC and lower expression of several nuclear-encoded mitochondrial pathways under OT. These opposing transcriptional patterns were associated with a stable genome-wide axis of transcriptional variation, suggesting that witnessing others’ suffering is associated with broader changes in monocyte gene expression involving distinct mitochondrial and nuclear-encoded gene programs.

## Results

### Established adversity-related transcriptional programs show condition-dependent differences

Participants completed a two-day, counterbalanced experimental protocol in which they viewed a 30-minute documentary depicting either animal trauma in the food industry (Observed Trauma; OT) or positive human-animal interactions (Observed Community; OC) during fMRI scanning (**Figure 1A**). Circulating monocytes were enriched from blood samples collected before and approximately 2 hours after each viewing session and profiled using bulk RNA sequencing. Within each condition, expression changes were quantified as the difference in log_2_-transformed counts per million (log_2_CPM) between post- and pre-viewing samples.

**Figure 1.**
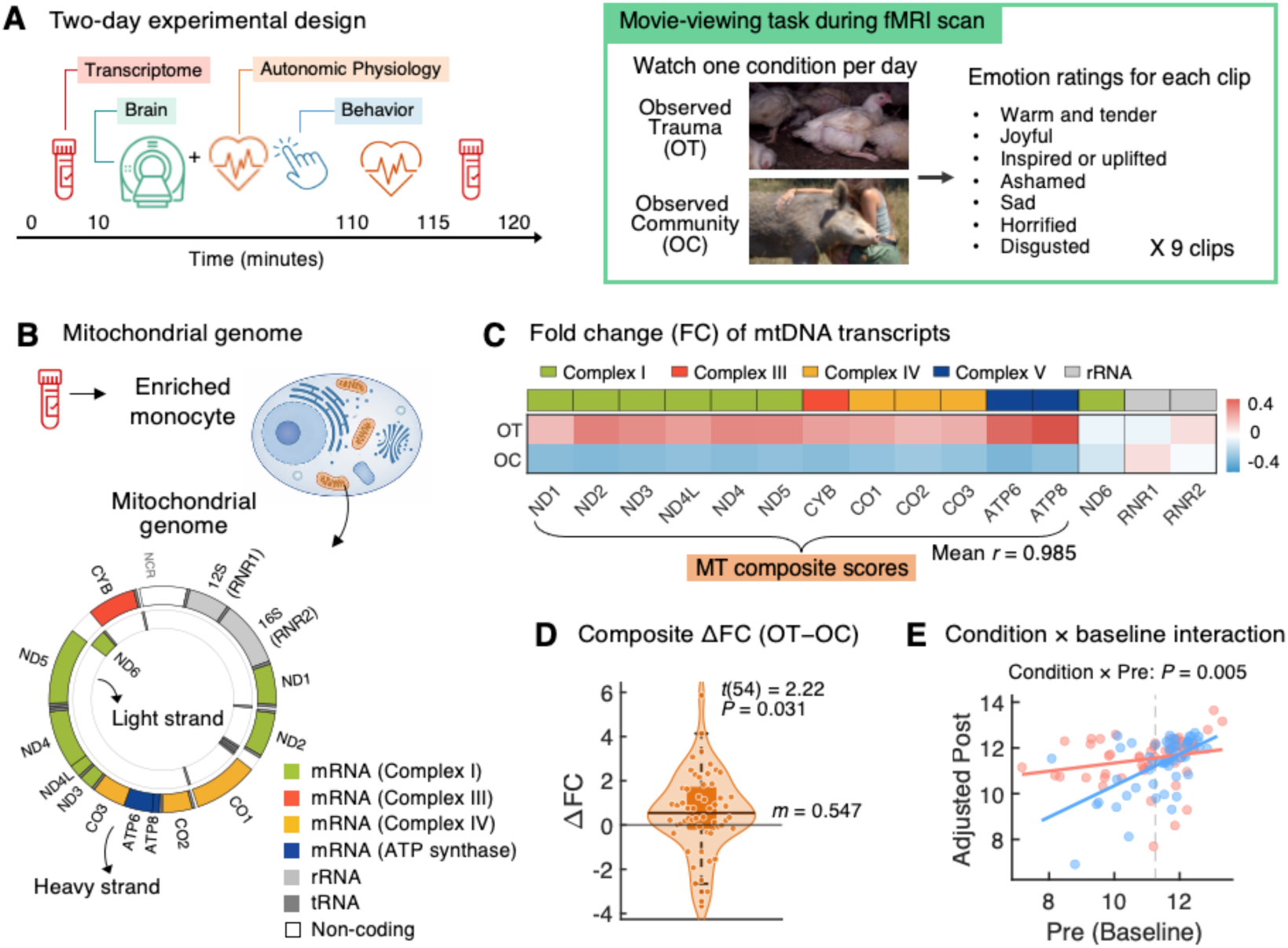
Divergent transcriptional dynamics of the mitochondrial genome. (A) Overview of the two-day experimental protocol. Each session began with an initial blood draw for baseline, an fMRI scan with intermittent self-reported emotional ratings for seven emotions during movie-viewing, autonomic physiological recording, and a post-viewing blood draw. Circulating monocytes were enriched from whole blood and analyzed using bulk RNA-sequencing. During the fMRI scan, participants watched a 30-minute animal documentary, which was either the Observed Trauma (OT) or Observed Community (OC) condition. Each participant watched one condition per day. (B) The human mitochondrial genome encodes 13 protein-coding mRNAs (Complex I, III, and IV, and ATP synthase), ribosomal RNAs, and transfer RNAs across the heavy (outer ring) and light (inner ring) strands. (C) Mean fold change (log_2_Post − log_2_Pre) in mtDNA-encoded transcript expression following OT (top) and OC (bottom). Red indicates increased expression and blue indicates decreased expression. Expression changes among the 12 heavy-strand mRNAs were highly correlated (mean pairwise *r* = 0.985). These transcripts were averaged to derive the MT composite score, excluding the light-strand transcript *MT-ND6*. (D) Differences in fold changes of the MT composite scores (OT−OC). Dots indicate each participant. The composite fold change was greater following OT than OC (*t*(54) = 2.22, *P* = 0.031). (E) Association between baseline and post-viewing MT composite expression under OT (red) and OC (blue). Post-viewing expression was adjusted for session order, sex, age, and participant-specific random intercepts using an LME model. Lines indicate condition-specific slopes, and the dashed vertical line marks the mean baseline expression. The baseline-post-viewing association was weaker under OT than OC (slope 0.18 vs 0.69; condition × baseline β = 0.25, *P* = 0.0051).

We first examined whether witnessing others’ suffering was associated with established adversity-related transcriptional patterns. We evaluated two gene sets derived from the conserved transcriptional response to adversity (CTRA): genes previously associated with high loneliness (LO; 61 genes retained after filtering) and genes associated with low loneliness (NL; 102 genes; **Figure S1A-B**). To determine whether post-viewing expression differed between conditions after accounting for baseline expression, we fitted a linear mixed-effects (LME) model predicting post-viewing expression from condition, mean-centered baseline expression, their interaction, and covariates (condition order, sex, and age), with a subject-specific random intercept (**Figure S1C)**. The NL gene set showed lower post-viewing expression under OT than OC (condition effect, *P* = 0.017), whereas the LO gene set showed no significant condition effect (*P* = 0.83). These findings suggest partial correspondence with established adversity-related transcriptional patterns, particularly in antiviral and antibody-related genes.

### Mitochondrial transcripts show a coordinated, condition-dependent expression pattern

We next examined mitochondrial transcriptional programs, given the central role of mitochondria in cellular adaptation to physiological demands. We focused first on 13 mitochondrial DNA (mtDNA)-encoded protein-coding transcripts (mt-mRNAs) spanning respiratory Complexes Ⅰ, Ⅲ, and Ⅳ as well as ATP synthase (**Figure 1B-C**). These transcripts accounted for 5.4-6.2% of mapped reads on average across conditions and time points. Their expression changes showed a consistent directional pattern, with mt-mRNAs generally increasing following OT and decreasing following OC. Among the 13 protein-coding transcripts, *MT-ND6* showed a markedly weaker response than the other mt-mRNAs. As the sole light-strand transcript, *MT-ND6* differs from the 12 heavy-strand transcripts in its transcriptional organization. Consistent with this distinction, changes in expression across the 12 heavy-strand transcripts were highly correlated (mean pairwise *r* = 0.985, 95% CI [0.978 0.992]), whereas correlations between *MT-ND6* and the other 12 transcripts were significantly lower (mean *r* = 0.461, 95% CI [0.197 0.669]; participant bootstrap, 1,000 resamples, *P* < 0.001). We therefore defined a mitochondrial transcript (MT) composite score as the mean log_2_CPM across the 12 heavy-strand protein-coding transcripts, excluding *MT-ND6*. This composite captures the coordinated expression pattern shared by the majority of the monocyte mt-mRNAs examined.

In an unadjusted within-subject comparison, the mean fold change in the MT composite was positive following OT (+0.290 ± 0.203 SE) and negative following OC (−0.258 ± 0.140 SE). The difference between conditions was significant (paired *t*(54) = 2.218, *P* = 0.031, Cohen’s *d* = 0.299), although the mean change within neither condition differed significantly from zero when tested separately (OT: *P* = 0.159; OC: *P* = 0.072; **Figure 1D**).

We then applied the same LME approach to the MT composite (**Supplementary Table 1**). The condition × baseline interaction was significant (β = 0.254, *SE* = 0.089, *t*(103) = 2.86, *P* = 0.0051; **Figure 1E**). Baseline expression was positively associated with post-viewing expression overall (β = 0.431, *SE* = 0.090, *P* = 5.0 × 10^-6^), but the strength of this association differed by condition. The estimated baseline slope was 0.686 under OC (*P* = 6.3 × 10^-6^), compared with 0.177 under OT (*P* = 0.095), indicating that baseline expression was strongly associated with subsequent expression under OC, whereas this relationship was substantially attenuated under OT. The condition main effect was not significant (β = −0.177, *SE* = 0.099, *P* = 0.077), although its negative estimate was directionally consistent with the higher fold change observed under OT than OC. At the individual-transcript level, the condition × baseline interaction was also significant after false discovery rate (FDR) correction for 12 of the 13 individual mt-mRNAs; *MT-ND6* was the exception (*P* = 0.589).

Together, these findings reveal a coordinated, condition-dependent pattern of mtDNA-encoded protein-coding transcript expression in circulating monocytes, characterized by divergent pre-to-post changes and a weaker association between baseline and post-viewing expression under OT.

### Nuclear-encoded mitochondrial pathways show condition-dependent expression patterns

The 13 mtDNA-encoded protein-coding transcripts examined above represent only a small fraction of the genes involved in mitochondrial function, with the vast majority encoded by the nuclear genome. To determine whether nuclear-encoded mitochondrial genes also exhibited condition-dependent expression patterns, we extended our analysis to 149 mitochondrial pathways defined by MitoCarta 3.0 (Rath et al., 2021). These pathways comprise 1,123 nuclear-encoded and 13 mtDNA-encoded genes organized into seven functional categories: metabolism, oxidative phosphorylation (OXPHOS), mitochondrial central dogma, small molecule transport, mitochondrial dynamics and surveillance, protein import, sorting, and homeostasis, and signaling (**Figure 2A**). Of the 149 pathways, 15 contain at least one mtDNA-encoded protein coding gene (mt-mRNA), all within the metabolism and OXPHOS categories. The remaining pathways comprised exclusively nuclear-encoded genes.

**Figure 2.**
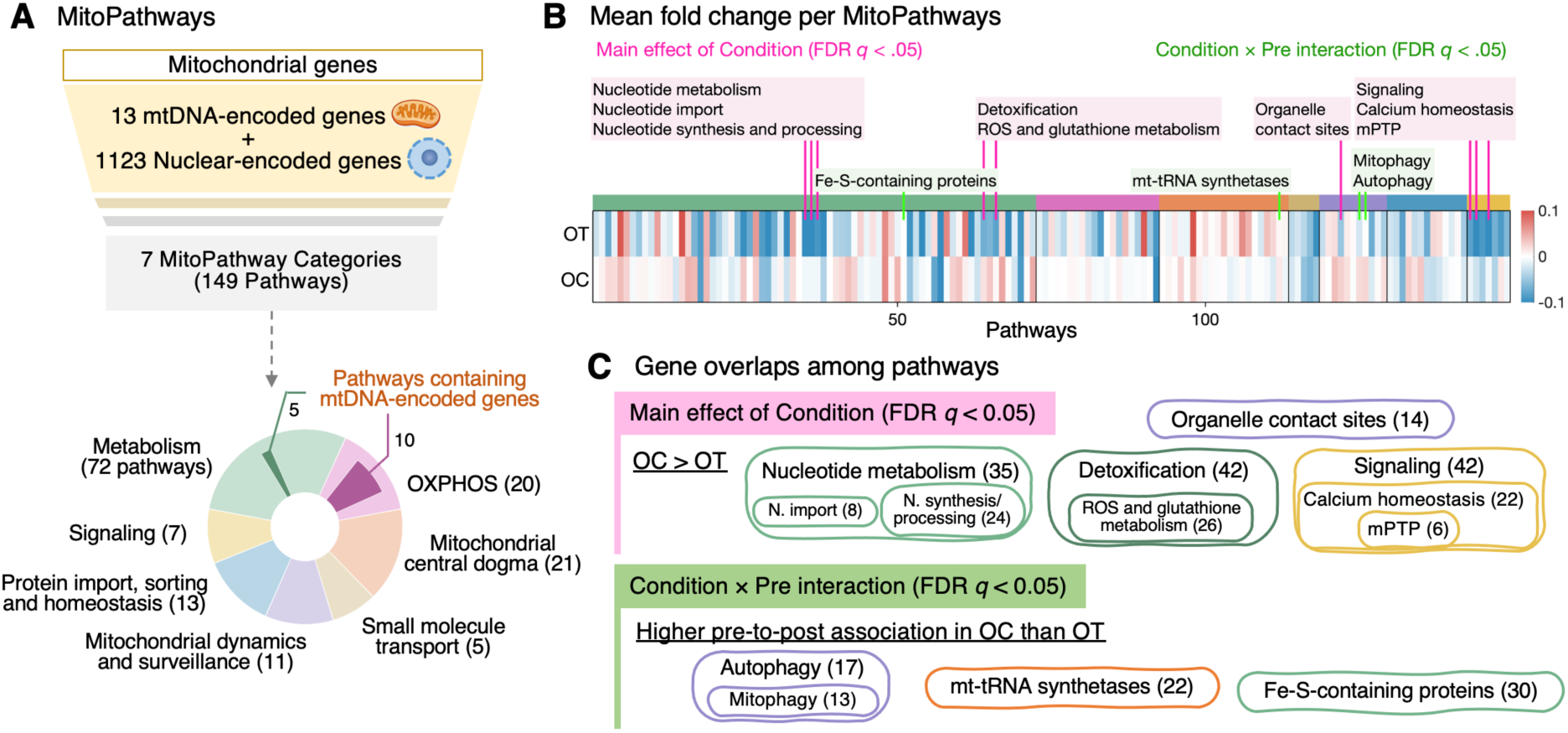
Nuclear-encoded mitochondrial pathways show condition-dependent expression. (A) MitoCarta 3.0 classifies 1,136 mitochondrial genes (13 mtDNA-encoded and 1,123 nuclear-encoded) into 149 pathways across seven functional categories. Ring segments indicate pathway counts; darker segments identify the 15 pathways containing mtDNA-encoded genes (5 metabolism, 10 OXPHOS). (B) Mean pathway fold change under OT (top) and OC (bottom), ordered by MitoCarta category (color bar). Pathway scores represent mean log_2_CPM across constituent genes. Vertical lines indicate significant condition main effects (magenta) or condition × baseline interactions (green) from LME models (*n* = 55, FDR *q* < 0.05 across 149 pathways). (C) Gene overlap among significant pathways. Nested outlines indicate pathways with shared gene sets; numbers indicate genes per pathway. None contained mtDNA-encoded genes.

Using the same LME modeling framework applied to the MT composite (post ∼ condition × pre + covariates + (1 | participant)), we identified 9 pathways with a significant main effect of condition and four with a significant condition × baseline interaction after false discovery rate (FDR) correction (*q* < 0.05; **Figure 2B**, **Table 1, Supplementary Table 2**). All 9 pathways with a significant condition main effect showed lower post-viewing expression under OT than OC (β = 0.032–0.091, FDR *q* = 0.008–0.028). In addition, the four pathways showing significant condition × baseline interactions exhibited weaker associations between baseline and post-viewing expression under OT than OC (β = 0.287–0.318, FDR *q* = 0.002–0.048), consistent with the baseline-dependent pattern observed for the MT composite. Notably, none of these 13 pathways contained mtDNA-encoded protein-coding genes, indicating that the observed effects were also present in exclusively nuclear-encoded mitochondrial gene sets.

**Table 1.** Condition-dependent expression of mitochondrial pathways. MitoCarta 3.0 pathways showing a significant condition main effect or condition × baseline interactions after Benjamini-Hochberg FDR correction (*q* < 0.05 across 149 pathways). Positive condition coefficients indicate higher post-viewing expression under OC at the mean baseline, whereas positive interaction coefficients indicate stronger baseline-post-viewing associations under OC. Asterisks indicate FDR-adjusted significance: \**q* < 0.05. \*\**q* < 0.01. \*\*\**q* < 0.001. Full results for all 149 pathways are provided in **Supplementary Table 2.**

| Category | MitoPathway | Condition |  |  |  | Baseline (pre) |  |  |  | Condition × Baseline |  |  |  |  |
|---|---|---|---|---|---|---|---|---|---|---|---|---|---|---|
|  |  | β | SE | *P* | *q* | β | SE | *P* | *q* | β | SE | *df* | *P* | *q* |
| Metabolism | Nucleotide metabolism | 0.041 | 0.010 | < .001 ** | 0.008 | 0.543 | 0.097 | < .001 *** | < .001 | 0.196 | 0.096 | 97.4 | 0.045 | 0.137 |
|  | Nucleotide import | 0.067 | 0.018 | < .001 * | 0.017 | 0.713 | 0.083 | < .001 *** | < .001 | -0.087 | 0.079 | 95.3 | 0.272 | 0.480 |
|  | Nucleotide synthesis and processing | 0.033 | 0.010 | 0.001 * | 0.024 | 0.541 | 0.104 | < .001 *** | < .001 | 0.166 | 0.104 | 96.4 | 0.113 | 0.268 |
|  | Fe-S-containing proteins | 0.006 | 0.008 | 0.495 | 0.698 | 0.419 | 0.088 | < .001 *** | < .001 | 0.287 | 0.087 | 103.0 | 0.001 * | 0.048 |
|  | Detoxification | 0.035 | 0.010 | 0.001 * | 0.024 | 0.481 | 0.092 | < .001 *** | < .001 | 0.213 | 0.092 | 97.6 | 0.023 | 0.099 |
|  | ROS and glutathione metabolism | 0.037 | 0.011 | 0.002 * | 0.028 | 0.493 | 0.083 | < .001 *** | < .001 | 0.218 | 0.084 | 97.6 | 0.011 | 0.087 |
| Mitochondrial central dogma | mt-tRNA synthetases | 0.005 | 0.008 | 0.533 | 0.742 | 0.562 | 0.086 | < .001 *** | < .001 | 0.303 | 0.081 | 99.8 | < .001 * | 0.015 |
| Mitochondrial dynamics and surveillance | Organelle contact sites | 0.032 | 0.008 | < .001 ** | 0.008 | 0.555 | 0.105 | < .001 *** | < .001 | 0.320 | 0.100 | 88.4 | 0.002 | 0.051 |
|  | Mitophagy | 0.021 | 0.010 | 0.042 | 0.219 | 0.402 | 0.068 | < .001 *** | < .001 | 0.318 | 0.072 | 101.5 | < .001 ** | 0.002 |
|  | Autophagy | 0.015 | 0.007 | 0.047 | 0.219 | 0.437 | 0.065 | < .001 *** | < .001 | 0.306 | 0.067 | 94.7 | < .001 ** | 0.002 |
| Signaling | Signaling | 0.033 | 0.008 | < .001 ** | 0.008 | 0.419 | 0.098 | < .001 *** | < .001 | 0.239 | 0.093 | 86.0 | 0.012 | 0.087 |
|  | Calcium homeostasis | 0.039 | 0.011 | < .001 * | 0.017 | 0.519 | 0.095 | < .001 *** | < .001 | -0.026 | 0.094 | 94.7 | 0.780 | 0.874 |
|  | Mitochondrial permeability transition pore | 0.091 | 0.022 | < .001 ** | 0.008 | 0.440 | 0.098 | < .001 *** | < .001 | 0.002 | 0.095 | 96.9 | 0.987 | 1.000 |

Several significant pathways shared overlapping genes and represented related biological processes (**Figure 2C**). Among the 9 pathways showing significant condition main effects, these overlaps were observed within four representative pathway groups: nucleotide metabolism, detoxification, organelle contact sites, and signaling. Similarly, the four pathways showing condition × baseline interactions included overlapping autophagy and mitophagy pathways, alongside mt-tRNA synthetases and Fe-S-containing proteins. Thus, the 13 significant pathways represented a smaller set of partially overlapping mitochondrial gene sets rather than entirely distinct sets of mitochondrial genes.

### Representative nuclear-encoded mitochondrial pathways exhibit distinct condition and baseline-dependent effects

To further characterize these effects, we examined representative pathways from the significant pathway sets (**Figure 3**). Among pathways showing significant condition main effects, we selected nucleotide metabolism, detoxification, organelle contact sites, and signaling, which encompassed the overlapping pathways identified in **Figure 2C**. All four showed lower post-viewing expression under OT than OC at the mean baseline level (nucleotide metabolism: β = 0.041, *P* < 0.001; detoxification: β = 0.035, *P* = 0.001; organelle contact sites: β = 0.032, *P* < 0.001; signaling: β = 0.033, *P* < 0.001). Baseline expression was comparable between conditions, whereas mean expression decreased from pre- to post-viewing under OT and remained relatively stable under OC (**Figure 3A**). This pattern contrasted with the MT composite, which showed a greater pre-to-post increase under OT than OC (Figure 1D), suggesting distinct condition-dependent expression responses between mtDNA-encoded transcripts and these nuclear-encoded mitochondrial pathways.

**Figure 3.**
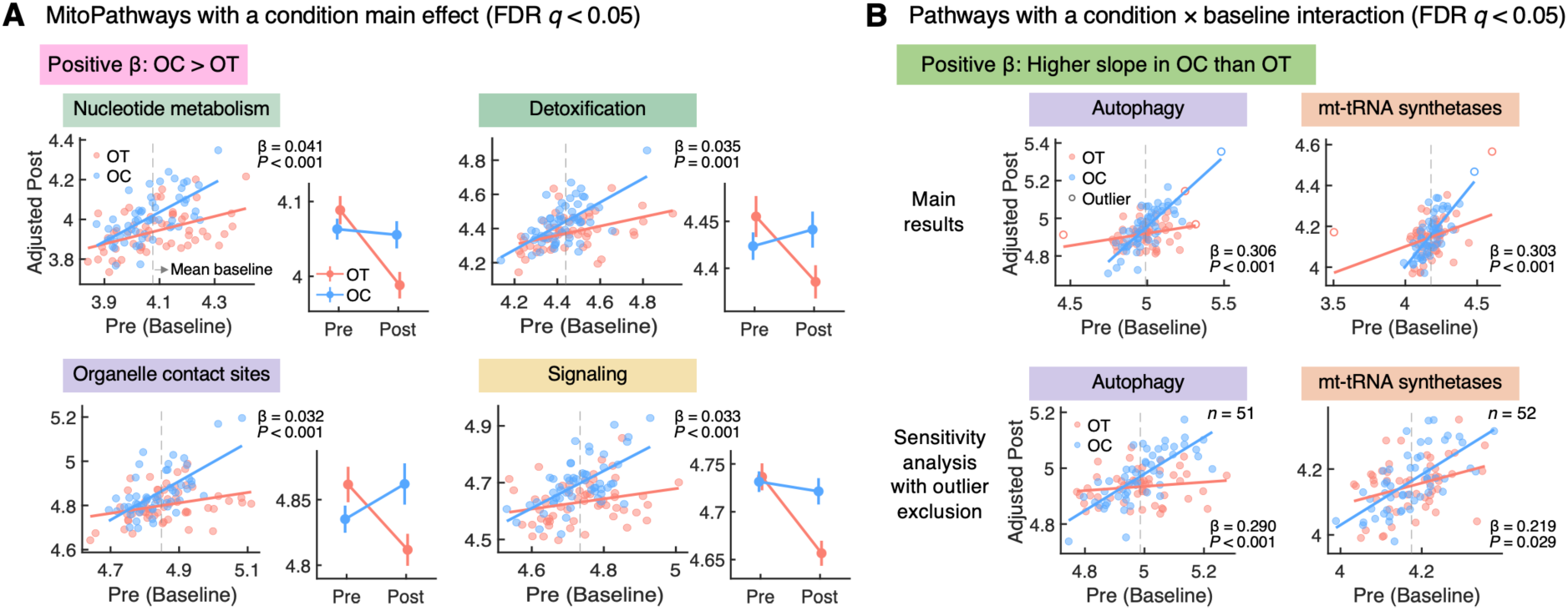
Representative mitochondrial pathways exhibit condition and baseline-dependent effects. (A) Representative pathways with significant condition main effects (FDR *q* < 0.05; Figure 2C). Left: post-viewing pathway scores versus baseline scores, adjusted for covariates and random intercepts per participant using LME models. Lines indicate condition-specific slopes; dashed lines mark mean baseline expression, at which condition effects are evaluated. Right: mean ± SE pathway scores before and after viewing. β and *P* indicate condition main effects from the LME model (OT = −1, OC = +1). All four pathways showed higher post-viewing expression under OC than OT at the mean baseline, contrasting with the higher MT composite fold change under OT (Figure 1D). (B) Representative pathways with significant condition × baseline interactions. Top: full sample (*n* = 55), with open circles indicating observations exceeding three scaled median absolute deviations from the condition-specific median in either baseline or post-viewing expression. Bottom: models refitted after excluding all observations from flagged participants (autophagy, *n* = 51; mt-tRNA synthetases, *n* = 52). β and *P* indicate interaction effects. Baseline-post-viewing associations were weaker under OT than OC in both pathways, with significant interactions persisting after outlier exclusion. OT, red; OC, blue.

Among pathways showing significant condition × baseline interactions, we focused on autophagy, which includes the mitophagy pathway (**Figure 2C**), mitochondrial tRNA synthetases (**Figure 3B**), and Fe-S-containing proteins (**Figure S2**). In all three pathways, baseline expression was strongly associated with post-viewing expression under OC, whereas this association was substantially weaker under OT (autophagy: β = 0.306, *P* < 0.001; slopes = 0.13 under OT and 0.74 under OC; mitochondrial tRNA synthetases: β = 0.303, *P* < 0.001; slopes = 0.26 and 0.87; Fe-S-containing proteins: β = 0.287, *P* = 0.001; slopes = 0.13 and 0.71). This pattern was consistent with the condition × baseline interaction observed for the MT composite (**Figure 1E**).

To assess the sensitivity of these interactions to extreme observations, we repeated the analyses after excluding outliers. Within each condition, observations with pre- or post-viewing expression values exceeding three scaled median absolute deviations from the condition-specific median were flagged (**Figure 3B**). All observations from participants with at least one flagged value were then excluded, and the models were refitted. The condition × baseline interaction remained significant for autophagy (*n* = 51, β = 0.290, *P* < 0.001) and mitochondrial tRNA synthetases (*n* = 52, β = 0.219, *P* < 0.001), but not for Fe-S-containing proteins (*n* = 51, β = 0.184, *P* = 0.072; **Figure S2**), which was therefore not considered further. The interaction effects also persisted when library sizes were recalculated after excluding mtDNA-derived reads, indicating that these findings were not explained by differences in the proportion of mitochondrial reads (**Supplementary Table 3**).

The condition main effects were similarly robust. All nine pathways with significant condition main effects in the primary analysis remained significant in the same direction after outlier exclusion (*P* = 0.0002–0.021) and recalculation of library sizes (*P* = 0.0002–0.004; **Supplementary Table 3**). Together, these analyses support robust condition effects in nuclear-encoded mitochondrial pathways (OT < OC), contrasting with the greater MT composite fold change under OT (OT > OC), alongside baseline-dependent effects in autophagy and mitochondrial tRNA synthetases that parallel those observed for the MT composite.

We further explored whether MT composite expression changes were associated with individual differences in personality traits, autonomic responses, or self-reported emotional experiences during movie viewing. We found no significant associations with animal attitudes, dispositional empathy, heart rate, skin conductance level, or any of seven emotional ratings (all *P* ≥ 0.068; **Supplementary Table 4**).

To assess whether condition-dependent transcriptional differences might reflect changes in monocyte subset composition, we examined canonical marker genes associated with classical (CD14, CCR2, SELL), intermediate (HLA-DRA), and non-classical (FCGR3A, CX3CR1, CDKN1C, LST1) monocytes. None of the eight markers showed a significant condition × time interaction (all *P* ≥ 0.163; **Supplementary Table 5**), and standardized effect sizes were small (∣*d*∣ ≤ 0.29). These findings provided no evidence of substantial condition-dependent changes in the expression of the selected monocyte subset markers.

### A stable genome-wide transcriptional axis is associated with mitochondrial transcript expression

To investigate whether the mitochondrial transcriptional patterns identified above were associated with broader genome-wide expression changes, we performed a principal component analysis (PCA) on the difference in fold change between conditions (ΔFC; OT minus OC) across 17,229 genes (**Figure 4A**). This approach identifies coordinated transcriptional patterns underlying interindividual variability in the OT−OC contrast. The first principal component (genePC1) explained 27.9% of the variance in ΔFC across participants (PC2, 11.2%, PC3, 7.8%) and showed high stability across participant bootstrap resamples (10,000 iterations, mean loading correlation *r* = 0.955, 95% CI [0.774, 0.990]). In contrast, subsequent components showed lower reproducibility (PC2: *r* = 0.734, 95% CI [0.041, 0.967]; PC3: *r* = 0.470, 95% CI [0.022, 0.988]). We therefore focus subsequent analyses on genePC1. Bootstrap ratios, calculated as the mean loading divided by its standard deviation across resamples, indicated that genes with larger absolute loadings also contributed more reliably to this axis (**Figure 4B**).

**Figure 4.**
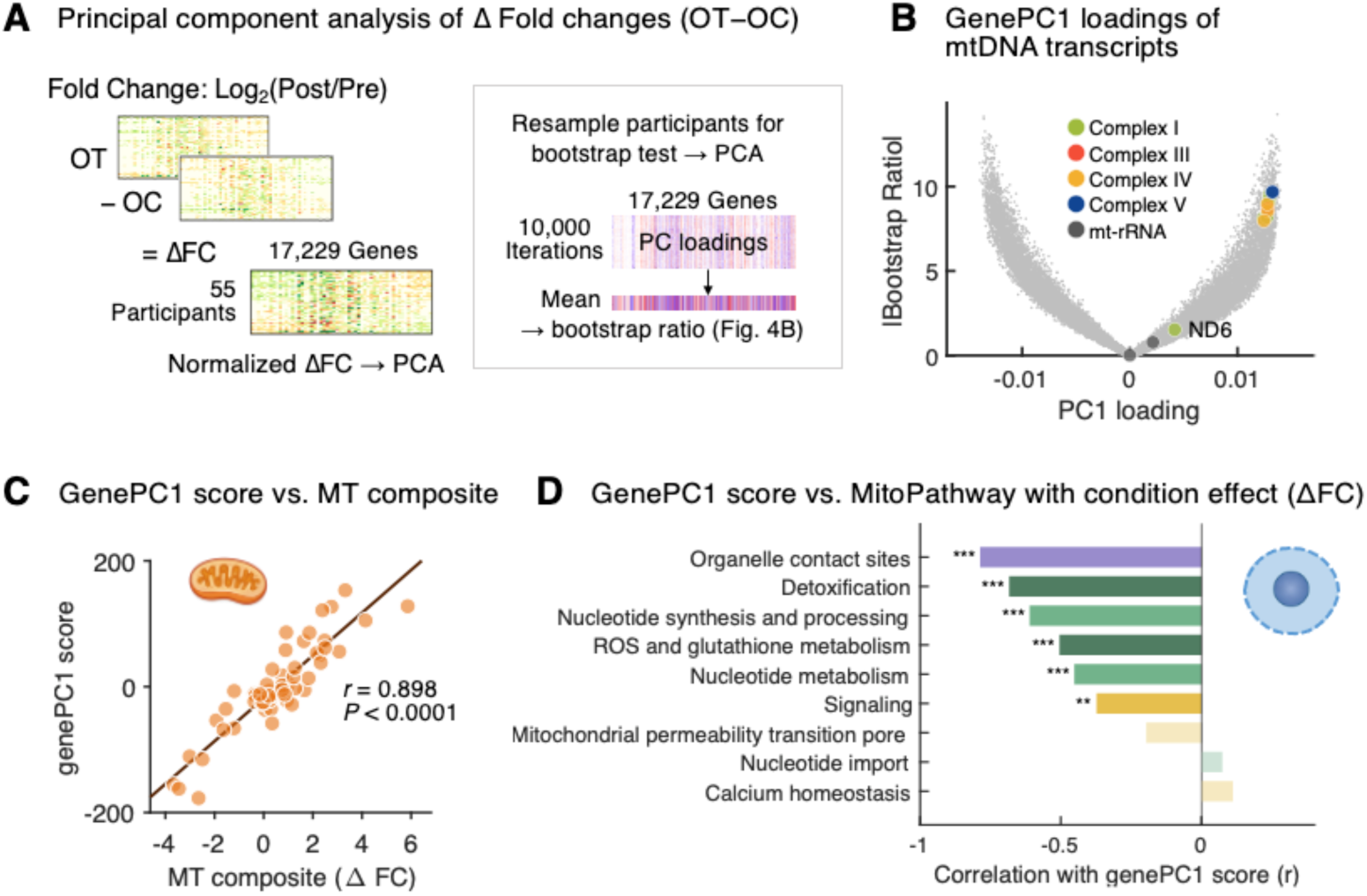
A genome-wide monocyte transcriptional axis tracks opposing changes in mtDNA-encoded transcripts and selected nuclear-encoded mitochondrial pathways. **(A)** Principal component analysis of the condition difference in fold change. For each participant and gene, fold change under OC was subtracted from that under OT (ΔFC), and the resulting 55 participants × 17,229 genes matrix was gene-wise z-scored and submitted to PCA. The first component (genePC1) explained 27.9% of the variance (PC2, 11.2%; PC3, 7.8%). To assess the reliability of each gene’s contribution, participants were resampled with replacement 10,000 times and the PCA was repeated; the bootstrap ratio is the mean genePC1 loading divided by its standard deviation across resamples. GenePC1 loadings were highly reproducible across resamples (mean loading correlation *r* = 0.955), unlike PC2 (*r* = 0.734) and PC3 (*r* = 0.470). (B) GenePC1 loading of each gene plotted against its absolute bootstrap ratio (grey). The 12 mt-mRNAs other than *MT-ND6* loaded positively and reliably on genePC1 and all ranked within the top 5% of genes (ranks 165–718 of 17,229; mean loading = 0.013), whereas *MT-ND6* and the mt-rRNAs lay near the center of the distribution. (C) Participants’ genePC1 scores plotted against the MT composite ΔFC (mean ΔFC of the 12 mt-mRNAs; Figure 1). The two were strongly correlated (Pearson *r* = 0.898, *P* < 0.0001; line, least-square fit), and this association persisted after excluding all mtDNA-encoded genes from PCA. (D) Pearson correlation between participants’ genePC1 scores and condition differences in fold change (ΔFC) for the nine mitochondrial pathways showing significant condition main effects (Figure 2C). Six pathways showed significant negative correlations with genePC1 (*r* = −0.373 to −0.787, *P* < 0.0001 to *P* = 0.0051), contrasting with the positive association observed for the MT composite in (C). Bars are colored by MitoCarta category; with distinct shades indicating detoxification and nucleotide metabolism subgroups within Metabolism. Faded bars indicate nonsignificant correlations (*P* > 0.05). Asterisks denote \*\**P* < 0.01, \*\*\**P* < 0.001.

Notably, the 12 mtDNA-encoded protein-coding transcripts (all except *MT-ND6*) showed strong positive loadings on genePC1, all ranking within the top 5% of 17,229 genes (**Figure 4B**). Consistent with these loadings, participants’ genePC1 scores were highly correlated with the condition difference in MT composite fold change (Pearson *r* = 0.898, *P* < 0.0001; **Figure 4C**). This correlation remained unchanged when all mtDNA-encoded genes were excluded before PCA (*r* = 0.898, *P* < 0.0001), indicating that the association between genePC1 and mitochondrial transcript expression was not solely attributable to the direct contribution of mtDNA-encoded genes to the component.

We next examined whether genePC1 was also associated with condition-dependent expression changes in the nuclear-encoded mitochondrial pathways identified above. In contrast to the strong positive association with the MT composite, genePC1 scores were negatively correlated with differential expression in several nuclear-encoded mitochondrial pathways (**Figure 4D**, **Supplementary Table 6**). The strongest correlations were observed for organelle contact sites (Pearson *r* = −0.787, *P* < 0.0001), detoxification (*r* = −0.684, *P* < 0.0001), and nucleotide synthesis and processing (*r* = −0.611, *P* < 0.0001). Additional negative associations were observed for ROS and glutathione metabolism, nucleotide metabolism, and signaling (*P* = 0.0001–0.0051). In contrast, mitochondrial permeability transition pore, nucleotide import, and calcium homeostasis showed no significant associations. These findings indicate that interindividual variation along genePC1 is associated with opposing expression changes in mtDNA-encoded transcripts and selected nuclear-encoded mitochondrial pathways.

To characterize the broader transcriptional programs associated with genePC1, we examined the 500 genes with the most positive and the 500 with the most negative loadings (**Figure S3**). Positively loaded genes showed increased expression following OT (mean FC = 0.151; one-sample *t*(54) = 2.42, *P* = 0.019) but not OC (mean FC = −0.016; *t*(54) = −0.30, *P* = 0.76), with a significant difference between conditions (paired *t*(54) = 2.07, *P* = 0.043). Conversely, negatively loaded genes showed decreased expression following OT (mean FC = −0.118; *t*(54) = −2.30, *P* = 0.025) but not OC (mean FC = 0.019; *t*(54) = 0.45, *P* = 0.65), with a significant difference between conditions (paired *t*(54) = −2.05, *P* = 0.045). Gene ontology enrichment analysis revealed distinct functional profiles at opposite ends of genePC1 (**Figure S4**). Positively loaded genes were enriched for macromolecule catabolism, proteolysis, metabolic regulation, and electron transport coupled proton transport (driven by mtDNA-encoded transcripts), broadly representing processes related to intracellular molecular turnover and metabolic regulation. In contrast, negatively loaded genes were enriched for vesicle-mediated transport, small GTPase-mediated signaling, and phosphorylation, representing processes involved in intracellular trafficking and signal transduction.

Finally, we examined whether genePC1 was aligned with the established adversity-related gene sets (CTRA) evaluated earlier. Both LO and NL gene sets were broadly distributed across the genePC1 loading spectrum, and neither showed a significant deviation in mean loading from randomly sampled gene sets of matched size (LO: *P* = 0.41; NL: *P* = 0.096; **Figure S1D-E**). Thus, although the lower expression of antiviral/antibody-related genes under OT partially corresponded with previously described adversity-related transcriptional patterns, neither CTRA gene set showed preferential representation at either end of genePC1. Together, these findings identify a stable genome-wide axis of transcriptional variation associated with opposing mitochondrial-related expression changes and broader functional differences between intracellular molecular regulation and cellular trafficking and signaling.

## Discussion

In this within-subject study, witnessing others’ suffering was associated with distinct transcriptional responses across mitochondrial and nuclear-encoded gene programs, embedded within broader variation in monocyte gene expression. Mitochondrial DNA-encoded protein-coding transcripts (mt-mRNAs, excluding *MT-ND6*) showed greater pre-to-post increase in expression following observed trauma (OT) than observed community (OC), whereas several nuclear-encoded mitochondrial pathways showed lower post-viewing expression under OT. These opposing patterns were also reflected in their associations with a stable genome-wide axis of transcriptional variation. In addition, the MT composite and selected nuclear-encoded pathways, including autophagy/mitophagy and mitochondrial tRNA synthetases, showed weaker associations between baseline and post-viewing expression under OT than OC. Together, these findings suggest that witnessing others’ suffering is associated with distinct mitochondrial-related transcriptional responses in circulating innate immune cells that extend beyond conventional inflammatory gene programs.

The divergent expression patterns of mt-mRNAs and nuclear-encoded mitochondrial pathways suggest that witnessing others’ suffering may engage distinct aspects of mitochondrial regulation. The 13 mtDNA-encoded protein-coding genes encode essential subunits of the oxidative phosphorylation (OXPHOS) machinery, which depends on coordinated contributions from the mitochondrial and nuclear genomes (*23*). Beyond energy production, mitochondria also function as signaling organelles that integrate metabolic and environmental information and communicate with other cellular compartments (*24*, *25*). These functions are particularly relevant to monocytes, whose metabolic regulation is closely linked to their immune functions (*26*). We observed a greater pre-to-post increase in mt-mRNA abundance following OT than OC, whereas several nuclear-encoded mitochondrial pathways showed lower expression under OT, including nucleotide metabolism, detoxification, organelle contact sites, and signaling. These pathways support distinct but interconnected aspects of mitochondrial biology, including nucleotide metabolism, redox homeostasis, inter-organelle communication, and intracellular signaling (*23*). Importantly, mitochondrial and nuclear mRNAs differ substantially in their production and turnover kinetics (*27*), providing a potential molecular context for their divergent expression patterns. Building on evidence that mitochondrial perturbations can engage nuclear transcriptional programs involved in cellular adaptation (*14*), these findings suggest that acute naturalistic psychosocial experiences may be associated with differential regulation of mitochondrial respiratory transcripts and nuclear-encoded programs supporting mitochondrial metabolism and homeostasis.

Beyond differences in average expression, mitochondrial-related transcriptional measures also showed condition-dependent associations between baseline and post-viewing expression. Previous studies have identified stable individual-specific patterns of peripheral blood gene expression (*28*) and substantial interindividual variation in the transcriptional responses of human monocytes to immune stimulation (*29*). In this study, baseline expression was strongly associated with post-viewing expression under OC, whereas this association was significantly attenuated under OT for the MT composite and nuclear-encoded pathways involving autophagy/mitophagy and mitochondrial tRNA synthetases. These findings suggest that transcriptional responses to observed suffering may depend not only on the exposure itself but also on the initial transcriptional state of circulating monocytes. Notably, our previous study using the same experimental paradigm identified baseline-dependent differences in pro-inflammatory CXCL8 expression following OT (*20*). Although the specific interaction patterns differed between CXCL8 and the mitochondrial-related measures, these findings collectively highlight the potential importance of pre-existing transcriptional states in shaping acute monocyte responses to psychosocial experiences.

The mitochondrial-related transcriptional differences were also embedded within broader genome-wide variation, as reflected by genePC1. This stable axis, derived from participant-level differences in the OT−OC contrast, was positively associated with MT composite changes and negatively associated with several nuclear-encoded mitochondrial pathways showing condition-dependent effects. Notably, its association with the MT composite persisted after excluding mtDNA-encoded genes from the PCA, indicating that the relationship extended beyond the direct contribution of mitochondrial transcripts. Gene ontology analysis revealed a functional contrast between genes associated with intracellular molecular turnover and metabolic regulation at the positive end of the axis and those involved in cellular trafficking and signaling at the negative end. This distinction is particularly relevant to monocytes, whose metabolic regulation is closely linked to immune functions (*26*), and whose trafficking and signaling machinery supports interactions with the surrounding environment (*30*). Together, these findings suggest that mitochondrial-related transcriptional responses to observed suffering were embedded within a broader organization of gene expression involving internal cellular regulation, and processes supporting environmental responsiveness.

These transcriptional patterns showed partial correspondence with the conserved transcriptional response to adversity (CTRA), which has been associated primarily with chronic or persistent social adversity (*6*). The pro-inflammatory gene set (LO) previously associated with loneliness showed no significant condition effect, whereas the antiviral- and antibody-related gene set (NL) showed lower post-viewing expression under OT than OC. However, neither gene set showed a preferential association with the positive or negative end of the genePC1 loading spectrum, suggesting that the broader transcriptional variation identified here was not dominated by canonical CTRA programs. Together with the mitochondrial and genome-wide findings, this partial correspondence suggests that acute psychosocial experiences may be associated with multiple dimensions of innate immune cell regulation beyond conventional inflammatory and antiviral transcriptional signatures. We also found no significant associations between MT composite expression changes and dispositional empathy, animal attitudes, autonomic responses, or self-reported emotional ratings during movie viewing. Thus, the mitochondrial-related transcriptional patterns were not readily captured by established adversity-related gene signatures or the psychological or autonomic measures examined. Nevertheless, the modest sample size and limited temporal resolution of these measurements leave open the possibility of smaller or temporally specific associations.

This study examined acute monocyte transcriptional responses to naturalistic psychosocial experiences within individuals, complementing previous social genomics work that has largely focused on chronic adversity and between-person differences. The counterbalanced, repeated-measures design, combined with multimodal physiological and behavioral assessments, provided an opportunity to investigate how witnessing others’ suffering is associated with short-term changes in peripheral immune cell gene expression. This paradigm examined responses to socially and morally salient information without directly exposing participants to physical threat or an experimentally imposed social evaluative stressor. The documentary films preserved the emotional and narrative complexity of witnessing animal suffering, providing a naturalistic context in which vicarious emotional engagement and peripheral immune responses could be examined together. However, this ecological richness also limits the ability to disentangle the contributions of negative affect, empathic concern, moral appraisal, and other features of the viewing experience. An important next step is therefore to determine which dimensions of observed suffering are most closely associated with the transcriptional patterns identified here.

Furthermore, although monocyte enrichment provided greater cellular specificity than mixed leukocyte samples, subset-resolved analyses will be necessary to establish whether these responses are shared across monocyte populations or concentrated in particular subsets. Importantly, changes in transcript abundance do not necessarily indicate corresponding changes in protein abundance or cellular function, and GO enrichment does not establish changes in pathway activity, mitochondrial respiration, or monocyte effector functions. Without direct measurements of monocyte mtDNA copy number or mitochondrial RNA turnover, the molecular processes underlying the observed mt-mRNA abundance differences remain unresolved. Studies with larger samples, more frequent sampling across a broader time course, and direct measurements of mitochondrial metabolism, protein abundance, and innate immune cell functions could further clarify the temporal dynamics and functional consequences of the observed mitonuclear transcriptional divergence.

Together, these findings demonstrate that witnessing others’ suffering is associated with acute changes in monocyte gene expression involving divergent mitochondrial and nuclear-encoded transcriptional programs. These transcriptional patterns were not fully captured by canonical adversity-related inflammatory signatures and encompassed broader variation in cellular metabolic, protein regulatory, and signaling processes. By demonstrating that exposure to socially and morally salient experiences without direct physical threat is associated with mitochondrial-related transcriptional changes in circulating monocytes, this study broadens our understanding of how vicarious trauma experiences relate to the molecular regulation of innate immune cells.

## Methods

### Study

This study was part of a larger multimodal investigation of observed trauma responses conducted at the Dartmouth Brain Imaging Center (DBIC), Dartmouth College, NH, USA, between February and November 2022, alongside functional Magnetic Resonance Imaging (fMRI), autonomic recording, and self-report measures, as previously described in (*20*). Although this dataset has been reported previously, the transcriptomic data have not been examined in relation to the present research questions. The present study focuses on peripheral immune transcriptomic data collected as part of this protocol. Participants visited the lab on two separate days for experimental sessions (mean interval = 10.09 ± 9.07 days). To reduce potential confounding by circadian variation, all sessions were conducted in the morning (8 AM-12 PM). The study protocol was approved by the Committee for the Protection of Human Subjects (Institutional Review Board) of Dartmouth College, and all participants provided written informed consent and received compensation for their participation. CTRA gene sets, mtDNA-encoded transcripts, and MitoPathway analyses were specified before analysis. The genome-wide PCA and the characterization of genePC1, including its relationship to mitochondrial and nuclear-encoded gene sets, were exploratory and conducted after the initial analyses.

### Participants

Fifty-five participants (age = 23.11 ± 5.76 years, 29 females) from the Upper Valley community in and around Hanover, NH, contributed to the present analyses. This subsample consists of participants from the previous study (*20*) with complete four-sample monocyte RNA-sequencing sets, yielding 220 samples. Of the 70 participants from whom blood was drawn, 15 were excluded for incomplete sampling or failure to meet RNA quality and sequencing criteria. Blood collection, sample processing, monocyte isolation, flow cytometry, and RNA sequencing procedures for this subsample have been detailed previously (*20*) and are briefly summarized below. A sensitivity power analysis (G*Power 3.1; F-test, linear multiple regression: fixed model, R^2^ increases) indicated that with *n* = 55, α = 0.05, and power = 0.80, the study was powered to detect interaction effects of f^2^ = 0.15, consistent with medium-sized effects.

### Self-report measurements

Participants responded to a set of self-report questionnaires during the pre-visit online survey. The personality measures include the Animal Attitude Scale 10-item version (AAS-10) (*31*) and the Interpersonal Reactivity Index (IRI) (*32*).

### Experimental tasks

Blood samples were collected before and after each fMRI session. Each fMRI session included multiple tasks, including a movie-viewing task. In the movie-viewing runs, participants watched one of two 30-minute animal documentaries, with condition order counterbalanced across participants. An observed trauma (OT) documentary depicted industrial farming, slaughter, and animal testing procedures, while an observed community (OC) documentary showed positive human-animal interactions, including animal rescue and affiliative contact with farm animals. Each documentary was presented across three runs and divided into nine clips, after each of which participants rated their emotional response along seven dimensions (warm and tender, joyful, inspired or uplifted, ashamed, sad, horrified, disgusted). Full task details are provided in (*20*). Documentary viewing began approximately 40 minutes after the baseline blood draw, and post-viewing samples were collected approximately 120 minutes after baseline.

### Blood collection, sample processing, and RNA sequencing

Peripheral blood was collected via venipuncture before and after each session (8 mL per draw, BD Vacutainer Mononuclear Cell Preparation Tubes with Sodium Heparin) and stored at 4 ℃ until processing. Plasma was separated by centrifugation, and mononuclear cells were washed before monocyte enrichment via negative selection (EasySep Human Monocyte Isolation Kit); enrichment quality was verified by flow cytometry (mean enrichment = 93.04% ± 9.88%). RNA was extracted from enriched monocytes (RNeasy Mini Kit) and submitted to the Genomics and Molecular Biology Shared Resource (GMBSR) at the Geisel School of Medicine at Dartmouth for library preparation and sequencing. Libraries were prepared using ribo-depletion (FastSelect HMR) followed by the Kapa RNA HyperPrep Kit and sequenced on a NextSeq2000 (Illumina; paired-end 50bp), targeting 30 million reads per sample. Reads were aligned to hg38 using STAR, and gene-level counts were quantified with featureCounts. All retained samples had more than 21 million mapped reads. Full details of these procedures are provided in (*20*).

### Expression quantification and normalization

Raw gene counts were processed in R v4.3.1 using the edgeR package (v.4.0.16; (*33*, *34*)). Lowly expressed genes were filtered using edgeR’s filterByExpr() function (grouped by condition), leaving 17,229 genes for downstream analysis. Library sizes were normalized using the Trimmed Mean of M values (TMM) method (calcNormFactors()), and normalized expression was converted to log_2_CPM using cpm(…, log = TRUE, prior.count = 0.25). To account for batch effects, log_2_CPM values were adjusted using ComBat (from the sva package 3.50.0), with sequencing batch specified as the batch variable and a model matrix including condition and visit (day 1 vs. day 2) to preserve biologically relevant variance while removing batch-associated variability. Condition-wise fold change was calculated as the post-minus pre-viewing difference in batch-corrected log_2_CPM for each gene.

### Conserved Transcriptional Response to Adversity (CTRA)

Conserved transcriptional response to adversity (CTRA) gene sets were derived from the leukocyte loneliness contrast of (*15*). They comprise transcripts over-expressed in high-lonely individuals (LO; 78 probes; pro-inflammatory) and in low-lonely individuals (NL; 123 probes; antiviral/antibody). Affymetrix HG-U133 Plus 2.0 probe identifiers were mapped to Ensembl gene identifiers with biomaRt (v2.58.2, affy_hg_u133_plus_2 → ensembl_gene_id, Homo sapiens GRCh38), yielding 87 and 169 unique genes; probes mapping to more than one gene contributed each mapped gene. Of these, 61 and 102, respectively, were present among the 17,229 filtered genes and were retained.

For each participant, condition, and time point, a composite score was computed as the mean log_2_CPM across the genes of each set. We fit a baseline-adjusted linear mixed-effects (LME) model for each composite score: post-viewing expression ∼ condition × baseline + order + sex + age + (1 | subject). Baseline was mean-centered so that condition effect is evaluated at the mean baseline. Condition was coded OT = −1, OC = +1. With effect coding, the condition coefficient represents half the adjusted OC−OT difference at the mean baseline, and the interaction coefficient represents half the difference between condition-specific baseline slopes. Models were fit by restricted maximum likelihood (REML), and degrees of freedom for fixed-effect tests were estimated using the Satterthwaite approximation. Analyses were performed in MATLAB R2024b (fitlme). This baseline-adjusted LME model was used for all subsequent expression analyses unless otherwise specified.

### Mitochondrial transcript analysis

All 13 mtDNA-encoded protein-coding transcripts (mt-mRNAs; *MT-ND1-6*, *MT-ND4L*, *MT-CO1-3*, *MT-ATP6*, *MT-ATP8*, *MT-CYB*) passed the expression filter and were analyzed. To quantify the contribution of mt-mRNAs to the transcriptome, we computed the proportion of each library mapping (mapped reads) to the 13 mt-mRNAs using the full unfiltered count data (60,617 genes × 220 samples).

Pairwise Pearson correlations of the condition difference in fold change (ΔFC; OT−OC) across participants were compared between the 12 heavy-strand mt-mRNAs and *MT-ND6*, the sole light-strand transcript, using participant-level bootstrap resampling (1,000 resamples). *MT-ND6* was therefore excluded from the MT composite score, defined as the mean log_2_CPM across the 12 heavy-strand transcripts. For the MT composite, we first compared fold change between conditions using a paired *t*-test.

To test whether post-viewing expression differed between conditions after accounting for baseline expression, we applied the baseline-adjusted LME model to each mt-mRNA and to the MT composite. P values for the 13 individual transcripts were corrected using the Benjamini-Hochberg (BH) FDR procedure (*q* < 0.05).

### MitoPathways

To characterize the nuclear- and mtDNA-encoded mitochondrial gene programs, we analyzed expression at the level of curated mitochondrial pathways defined in MitoCarta 3.0 (*23*). Pathway gene memberships were taken from the MitoPathways ontology and matched to our data by Ensembl identifier, yielding 149 pathways with members present in the filtered expression data. These pathways span seven top-level categories: metabolism, OXPHOS, mitochondrial central dogma, small molecule transport, mitochondrial dynamics and surveillance, protein import/sorting/homeostasis, and signaling. For each pathway and sample, a pathway score was computed as the mean log_2_CPM of its member genes. A pathway was annotated as mtDNA-containing if at least one member gene was an mt-mRNA.

Pathway scores were analyzed with the same linear mixed-effects model and coding used for MT composite: post ∼ condition × baseline + order + sex + age + (1 | subject). P values were corrected across the 149 pathways using the BH FDR procedure (*q* < 0.05).

We ran two sensitivity analyses for pathways significant in the primary analysis. First, outlier exclusion: within each condition, an observation was flagged if its pre- or post-viewing score deviated from the condition-specific median by more than three scaled median absolute deviations. All four observations from flagged participants were then removed for that pathway, and the model was refitted. Second, mitochondrial read share: mtDNA-encoded reads make up a variable fraction of each library, so they can shift the log_2_CPM values of nuclear genes. To assess the sensitivity of the results to differences in mitochondrial read abundance, we recalculated library sizes and TMM normalization factors after excluding the 37 mtDNA-encoded genes, repeated the ComBat correction, and refitted the models.

### Associations with individual-difference measures

To examine whether MT composite responses were associated with individual differences, we computed Pearson correlations between MT composite fold change and personality traits, autonomic responses, and emotion ratings. Personality traits comprised the Animal Attitude Scale and the Interpersonal Reactivity Index. For the autonomic responses, heart rate (HR) and skin conductance level (SCL) were recorded during movie-viewing and each averaged within conditions. Emotion ratings comprised the seven emotion ratings, each averaged across the nine clips within conditions. For condition-varying measures, ΔFC was correlated with the corresponding OT−OC difference in the measure, and FC was correlated with the measure within each condition separately. Personality traits were measured once and were therefore correlated with ΔFC only. Participants with poor-quality physiological recordings, identified as in (*20*), were excluded from the HR (*n* = 53) and SCL (*n* = 49) analyses.

### Monocyte subset markers

To assess whether condition effects might reflect shifts in monocyte subset composition, we examined canonical markers of classical (*CD14*, *CCR2*, *SELL*), intermediate (*HLA-DRA*), and non-classical (*FCGR3A*, *CX3CR1*, *CDKN1C*, *LST1*) monocytes. Expression was quantified as log2 transcripts per million (TPM), and each marker was analyzed with an LME model of the form expression ∼ condition × time + order + batch + sex + age + (1 | subject). The condition × time interaction was the term of interest. Effect sizes were expressed as Cohen’s d for the within-participant difference in pre-to-post change (OT−OC), with 95% confidence intervals estimated by participant-level bootstrap resampling (5,000 resamples).

### Principal Component Analysis

For each participant and gene, we computed the condition difference in fold change (ΔFC; OT−OC). The resulting 55 participants × 17,229 genes matrix was z-scored within each gene and submitted to PCA, with participants as observations and genes as variables. Each component yielded a gene-wise loading vector and a participant-wise score. Higher genePC1 scores indicate stronger expression of the genePC1 pattern in the OT−OC contrast.

To assess component stability, participants were resampled with replacement 10,000 times and the PCA was recomputed. Bootstrapped loadings were sign-aligned to the original solution, and stability was quantified as the Pearson correlation between the original and bootstrapped loading vectors. The reliability of each gene’s contribution to genePC1 was quantified with a bootstrap ratio: the mean loading across resamples divided by its standard deviation.

To test whether the association between genePC1 and the MT composite reflected the direct contribution of mtDNA-encoded genes, we repeated the PCA after excluding all mtDNA-encoded genes that passed filtering (*n* = 34). We then correlated the resulting participant scores with the MT composite ΔFC. Associations between genePC1 scores and the ΔFC of the nine MitoPathways with significant condition effects were tested with Pearson correlations.

For exploratory characterization, we selected the 500 genes with the most positive and the 500 with the most negative genePC1 loadings. All 1,000 genes had ∣bootstrap ratio∣ > 5. For each set, mean FC across genes was compared against zero within each condition (one-sample *t*-tests) and between conditions (paired *t*-tests).

To test whether the CTRA sets were represented along genePC1, each set’s mean genePC1 loading was compared against a null distribution (two-sided). The null consisted of 10,000 randomly drawn gene sets of the same size from the same pool of 17,229 genes.

### Gene Ontology Enrichment analysis

Functional enrichment analysis was performed separately on the 500 genes with the most positive and the 500 with the most negative genePC1 loadings using ShinyGO 0.85.1. To avoid tissue expression bias, enrichment was tested against a custom background of all 17,229 expressed genes included in the PCA. Over-representation of Gene Ontology Biological Process (GO-BP) terms was assessed with BH FDR correction. Terms with FDR (*q* < 0.05) and at least three genes were considered significant.

Because significant GO-BP terms are frequently nested within the GO hierarchy and thus highly redundant, we reduced them to representative categories using rrvgo v1.14.2. Semantic similarity between terms was computed on the BP ontology using the relevance (Rel) measure, and terms were grouped by hierarchical clustering at a similarity threshold of 0.7. Within each group, the term with the strongest enrichment (lowest FDR) was designated as the representative term.

## Acknowledgments

**Funding**

The work was supported by NIMH R01 MH116026 (T.D.W.), R37 MH076136 (T.D.W.), NIBIB R01 EB026549 (T.D.W.), Animals Australia for a Kinder World (T.D.W.), and NIH R35 HL155458 (C.V.J.). Bulk RNA sequencing was carried out in the Genomics and Molecular Biology Shared Resource (RRID:SCR_021293) at Dartmouth, which is supported by NCI Cancer Center Support Grant 5P30CA023108 and NIH S10 (1S10OD030242) awards.

This manuscript was proofread and checked for grammar and typo using ChatGPT and Claude.

## Author contributions

Conceptualization: B.K.L., M.C.K., C.V.J., T.D.W.; Data curation: B.K.L., M.C.K.; Formal analysis: B.K.L., X.L., M.P., C.V.J., T.D.W.; Funding acquisition: F.W.K., C.V.J., T.D.W.; Investigation: B.K.L., M.C.K., F.W.K., X.L.; Project administration: B.K.L., T.D.W.; Resources: B.K.L., F.W.K., X.L., C.V.J.; Supervision: T.D.W.; Visualization: B.K.L.; Writing – original draft: B.K.L., T.D.W.; Writing – review & editing: B.K.L., M.P., C.V.J., T.D.W.

## Data, code, and materials availability

All data needed to evaluate the conclusions in the paper are present in the paper and/or the Supplementary Materials. Processed gene-level count matrices have been deposited in the NCBI Gene Expression Omnibus under accession GSE352855 (reviewer access token provided with this submission; the record will be released on publication). Preprocessed data supporting the findings and figures of this study are available on Zenodo at https://doi.org/10.5281/zenodo.23681460. All figures from the main experimental cohort in the main manuscript and the supplementary materials can be reproduced from these data and code. Raw sequencing reads are not publicly deposited to protect participant privacy and comply with the ethical restrictions approved by the Institutional Review Board (IRB) of Dartmouth College. These data are available from the first author (B.K.L.) upon request and requests will be answered within 30 days.

## References

1. F. B. M. De Waal, Putting the Altruism Back into Altruism: The Evolution of Empathy. Annu. Rev. Psychol. 59, 279–300 (2008).

2. M. Masalha, M. Cohen, M. Aloni, E. Toledo, E. Stahl, D. Shachar, O. Eliav, S. Fuchs, S. Atzil, Social Physiology: Human homeostasis is more efficient in social proximity. Sci. Adv. 12, eaeg4937 (2026).

3. J. T. Cacioppo, Social neuroscience: Autonomic, neuroendocrine, and immune responses to stress. Psychophysiology 31, 113–128 (1994).

4. V. Engert, J. Smallwood, T. Singer, Mind your thoughts: Associations between self-generated thoughts and stress-induced and baseline levels of cortisol and alpha-amylase. Biol. Psychol. 103, 283–291 (2014).

5. V. Engert, R. Linz, J. A. Grant, Embodied stress: The physiological resonance of psychosocial stress. Psychoneuroendocrinology 105, 138–146 (2019).

6. S. W. Cole, The Conserved Transcriptional Response to Adversity. Curr. Opin. Behav. Sci. 28, 31–37 (2019).

7. G. E. Hodes, M. L. Pfau, M. Leboeuf, S. A. Golden, D. J. Christoffel, D. Bregman, N. Rebusi, M. Heshmati, H. Aleyasin, B. L. Warren, B. Labonté, S. Horn, K. A. Lapidus, V. Stelzhammer, E. H. F. Wong, S. Bahn, V. Krishnan, C. A. Bolaños-Guzman, J. W. Murrough, M. Merad, S. J. Russo, Individual differences in the peripheral immune system promote resilience versus susceptibility to social stress. Proc. Natl. Acad. Sci. 111, 16136–16141 (2014).

8. O. K. Sial, B. L. Warren, L. F. Alcantara, E. M. Parise, C. A. Bolaños-Guzmán, Vicarious social defeat stress: Bridging the gap between physical and emotional stress. J. Neurosci. Methods 258, 94–103 (2016).

9. M. R. Irwin, S. W. Cole, Reciprocal regulation of the neural and innate immune systems. Nat. Rev. Immunol. 11, 625–632 (2011).

10. B. K. Chacko, P. A. Kramer, S. Ravi, M. S. Johnson, R. W. Hardy, S. W. Ballinger, V. M. Darley-Usmar, Methods for defining distinct bioenergetic profiles in platelets, lymphocytes, monocytes, and neutrophils, and the oxidative burst from human blood. Lab. Invest. 93, 690–700 (2013).

11. N. Bobba-Alves, R.-P. Juster, M. Picard, The energetic cost of allostasis and allostatic load. Psychoneuroendocrinology 146, 105951 (2022).

12. G. Barshad, A. Blumberg, T. Cohen, D. Mishmar, Human primitive brain displays negative mitochondrial-nuclear expression correlation of respiratory genes. Genome Res., doi: 10.1101/gr.226324.117 (2018).

13. N. J. Kramer, G. Prakash, R. S. Isaac, K. Choquet, I. Soto, B. Petrova, H. E. Merens, N. Kanarek, L. S. Churchman, Regulators of mitonuclear balance link mitochondrial metabolism to mtDNA expression. Nat. Cell Biol. 25, 1575–1589 (2023).

14. P. M. Quirós, A. Mottis, J. Auwerx, Mitonuclear communication in homeostasis and stress. Nat. Rev. Mol. Cell Biol. 17, 213–226 (2016).

15. S. W. Cole, L. C. Hawkley, J. M. Arevalo, C. Y. Sung, R. M. Rose, J. T. Cacioppo, Social regulation of gene expression in human leukocytes. Genome Biol. 8, R189 (2007).

16. S. W. Cole, L. C. Hawkley, J. M. G. Arevalo, J. T. Cacioppo, Transcript origin analysis identifies antigen-presenting cells as primary targets of socially regulated gene expression in leukocytes. Proc. Natl. Acad. Sci. 108, 3080–3085 (2011).

17. S. W. Cole, J. P. Capitanio, K. Chun, J. M. G. Arevalo, J. Ma, J. T. Cacioppo, Myeloid differentiation architecture of leukocyte transcriptome dynamics in perceived social isolation. Proc. Natl. Acad. Sci. 112, 15142–15147 (2015).

18. V. Engert, F. Plessow, R. Miller, C. Kirschbaum, T. Singer, Cortisol increase in empathic stress is modulated by emotional closeness and observation modality. Psychoneuroendocrinology 45, 192–201 (2014).

19. S. Keum, H.-S. Shin, Neural Basis of Observational Fear Learning: A Potential Model of Affective Empathy. Neuron 104, 78–86 (2019).

20. B. K. Lux, M. C. Kos, D. Ward, F. W. Kolling, X. Li, C. V. Jakubzick, T. D. Wager, Vicarious trauma primes innate immunity and reconfigures human brain networks. bioRxiv, 2026.01.05.697785 (2026).

21. S. S. Dickerson, T. L. Gruenewald, M. E. Kemeny, When the Social Self Is Threatened: Shame, Physiology, and Health. J. Pers. 72, 1191–1216 (2004).

22. S. S. Dickerson, M. E. Kemeny, N. Aziz, K. H. Kim, J. L. Fahey, Immunological Effects of Induced Shame and Guilt. Psychosom. Med. 66, 124–131 (2004).

23. S. Rath, R. Sharma, R. Gupta, T. Ast, C. Chan, T. J. Durham, R. P. Goodman, Z. Grabarek, M. E. Haas, W. H. W. Hung, P. R. Joshi, A. A. Jourdain, S. H. Kim, A. V. Kotrys, S. S. Lam, J. G. McCoy, J. D. Meisel, M. Miranda, A. Panda, A. Patgiri, R. Rogers, S. Sadre, H. Shah, O. S. Skinner, T.-L. To, M. A. Walker, H. Wang, P. S. Ward, J. Wengrod, C.-C. Yuan, S. E. Calvo, V. K. Mootha, MitoCarta3.0: an updated mitochondrial proteome now with sub-organelle localization and pathway annotations. Nucleic Acids Res. 49, D1541–D1547 (2021).

24. V. Eisner, M. Picard, G. Hajnóczky, Mitochondrial dynamics in adaptive and maladaptive cellular stress responses. Nat. Cell Biol. 20, 755–765 (2018).

25. M. Picard, O. S. Shirihai, Mitochondrial signal transduction. Cell Metab. 34, 1620–1653 (2022).

26. L. A. J. O’Neill, R. J. Kishton, J. Rathmell, A guide to immunometabolism for immunologists. Nat. Rev. Immunol. 16, 553–565 (2016).

27. E. McShane, M. Couvillion, R. Ietswaart, G. Prakash, B. M. Smalec, I. Soto, A. R. Baxter-Koenigs, K. Choquet, L. S. Churchman, A kinetic dichotomy between mitochondrial and nuclear gene expression processes. Mol. Cell 84, 1541–1555.e11 (2024).

28. J. P. Radich, M. Mao, S. Stepaniants, M. Biery, J. Castle, T. Ward, G. Schimmack, S. Kobayashi, M. Carleton, J. Lampe, P. S. Linsley, Individual-specific variation of gene expression in peripheral blood leukocytes. Genomics 83, 980–988 (2004).

29. B. P. Fairfax, P. Humburg, S. Makino, V. Naranbhai, D. Wong, E. Lau, L. Jostins, K. Plant, R. Andrews, C. McGee, J. C. Knight, Innate Immune Activity Conditions the Effect of Regulatory Variants upon Monocyte Gene Expression. Science 343, 1246949 (2014).

30. C. Shi, E. G. Pamer, Monocyte recruitment during infection and inflammation. Nat. Rev. Immunol. 11, 762–774 (2011).

31. H. Herzog, S. Grayson, D. McCord, Brief Measures of the Animal Attitude Scale. Anthrozoös 28, 145– 152 (2015).

32. M. H. Davis, Interpersonal reactivity index. (1980).

33. M. D. Robinson, D. J. McCarthy, G. K. Smyth, edgeR: a Bioconductor package for differential expression analysis of digital gene expression data. Bioinformatics 26, 139–140 (2010).

34. M. D. Robinson, A. Oshlack, A scaling normalization method for differential expression analysis of RNA-seq data. Genome Biol. 11, R25 (2010).

