## Supplementary Materials for "Witnessing others’ suffering is associated with divergent mitochondrial transcriptional programs in human monocytes"

**This PDF file includes:**

Figures S1 to S4

Tables S1 to S6

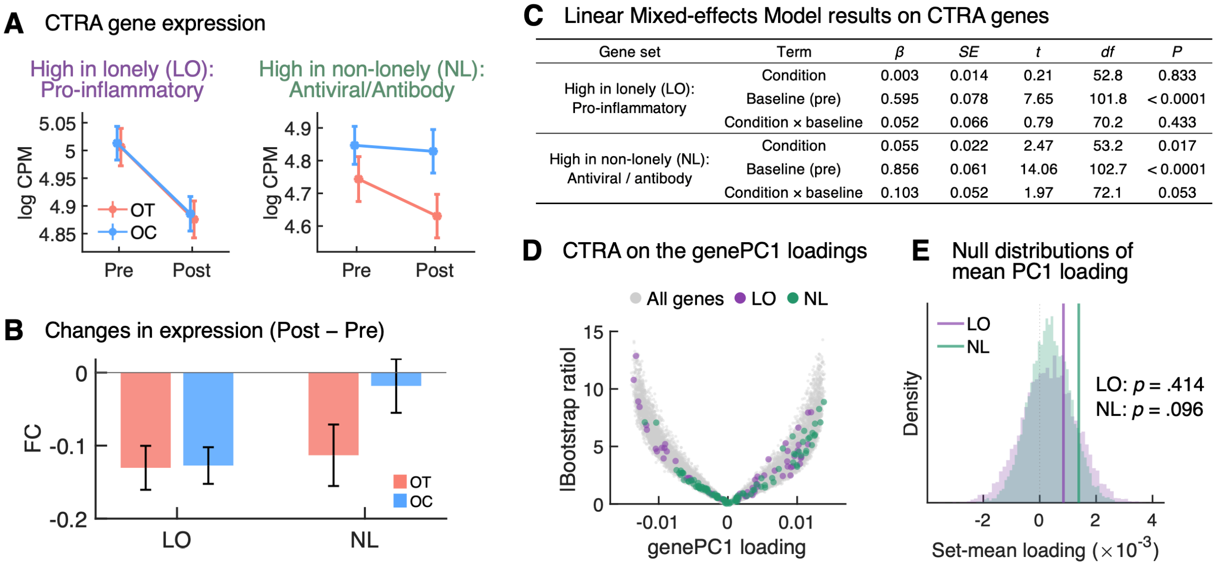


| Gene set | Term | β | SE | t | df | P |
|---|---|---|---|---|---|---|
| High in lonely (LO): Pro-inflammatory | Condition | 0.003 | 0.014 | 0.21 | 52.8 | 0.833 |
| | Baseline (pre) | 0.595 | 0.078 | 7.65 | 101.8 | < 0.0001 |
| | Condition × baseline | 0.052 | 0.066 | 0.79 | 70.2 | 0.433 |
| High in non-lonely (NL): Antiviral / antibody | Condition | 0.055 | 0.022 | 2.47 | 53.2 | 0.017 |
| | Baseline (pre) | 0.856 | 0.061 | 14.06 | 102.7 | < 0.0001 |
| | Condition × baseline | 0.103 | 0.052 | 1.97 | 72.1 | 0.053 |

**Figure S1. The CTRA gene sets are separable from the genePC1 axis.** Two a priori conserved transcriptional response to adversity (CTRA) gene sets were scored. Genes up-regulated in lonely individuals (pro-inflammatory, LO) and genes up-regulated in non-lonely individuals (antiviral/antibody, NL). Matching ENSEMBL identifiers against the filtered expression matrix recovered 61 of 87 LO genes and 102 of 169 NL genes. Each participant's score is the mean $\log_2$CPM across the genes of a set, computed for each condition and time point. (A) Pre- to post-viewing change of each composite by condition (red, OT; blue, OC; mean ± SEM, $n$ = 55). (B) Fold change in each set (mean ± SEM). (C) LME model results for each set (model as in Figure 1E). The NL set showed lower expression after OT than after OC (condition effect, $P$ = 0.017), whereas the LO set showed no condition or condition × baseline effect. (D) Position of the CTRA genes on the genePC1 loading × absolute bootstrap ratio map. Purple, LO genes; green, NL genes. (E) Set-level permutation test on the mean genePC1 loading. For each set, the observed mean loading (vertical line) was compared against 10,000 random gene sets of matched size drawn from the same 17,229 gene pool, with a two-sided $P$ on the signed mean. Both observed means fall within their null distributions (LO: mean loading = 0.00084, null $SD$ = 0.00098, $P$ = 0.414; NL: 0.00138, 0.00075, $P$ = 0.096, two-sided). The NL null is narrower because it averages over more genes (102 vs. 61).

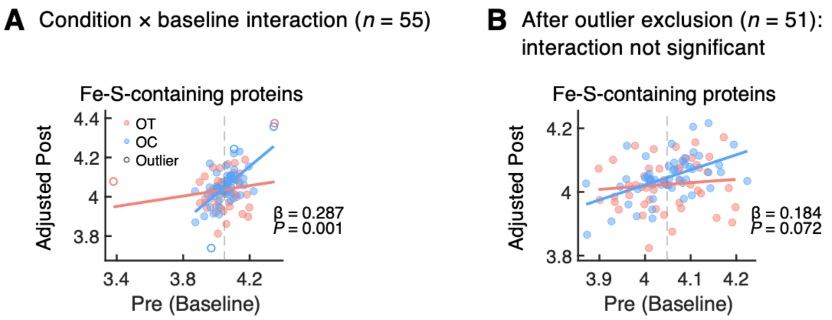


**Figure S2. Condition × baseline interaction for Fe-S-containing proteins.** (A) Post-viewing versus baseline scores, adjusted for condition order, sex, age, and participant random intercept, for all participants ($n$ = 55). Lines show the model-estimated condition-specific slopes, and the dashed line marks the mean baseline. Open circles mark observations whose pre- or post-viewing score deviated from the condition median by more than three scaled median absolute deviations. The interaction was significant after FDR correction across 149 pathways (β = 0.287, $P$ = 0.001, **Table S3**). (B) The same model refitted after removing all samples from the four participants with flagged observations ($n$ = 51). The interaction was no longer significant (β = 0.184, $P$ = 0.072). OT, red; OC, blue.

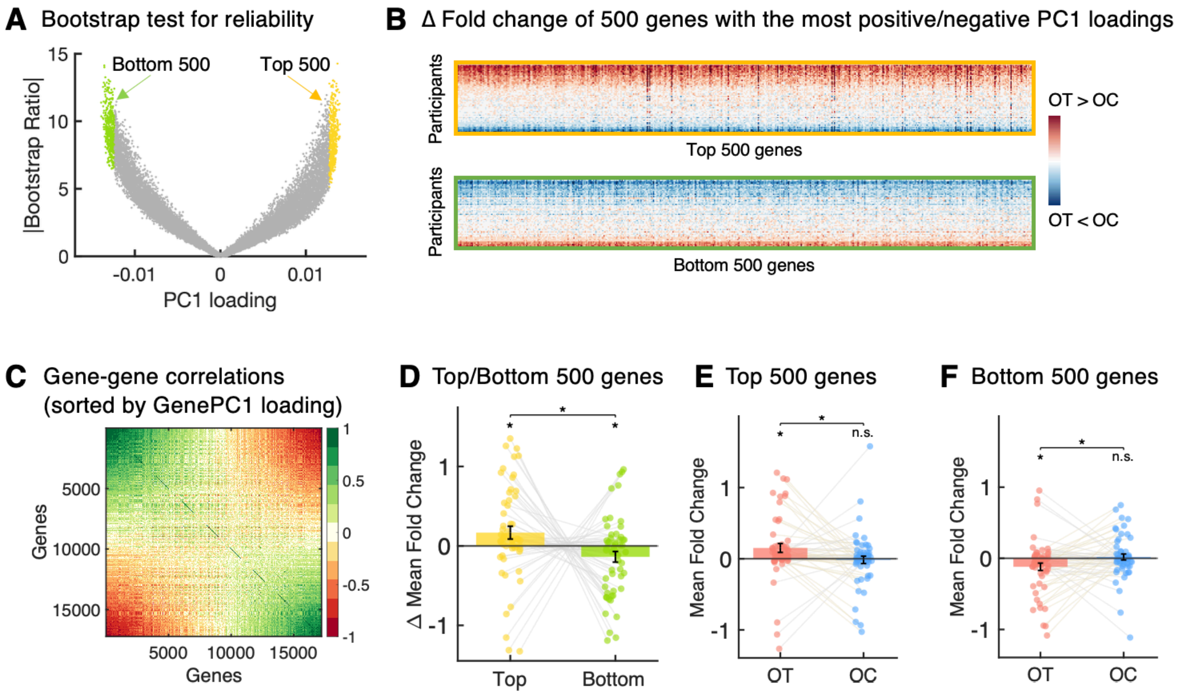


**Figure S3. Principal component analysis of genome-wide monocyte transcription.** (A) GenePC1 loading of 17,229 genes (x-axis) plotted against their absolute bootstrap ratios (y-axis; mean loading divided by its standard deviation across 10,000 bootstrap resamples of participants). The 500 genes with the most positive loadings (top 500, yellow) and the 500 with the most negative loadings (bottom 500, green) are highlighted; all have |bootstrap ratio| > 5. (B) Difference in fold change between conditions ($\Delta FC = FC_{OT} - FC_{OC}$; $FC = \log_2 Post - \log_2 Pre$) for each participant (rows) across the top 500 (top) and bottom 500 (bottom) genes (columns). Participants are ordered by their mean ΔFC across the top 500 genes. Red indicates ΔFC > 0 (OT > OC) and blue indicates ΔFC < 0 (OT < OC). (C) Pairwise correlations between all 17,229 genes across 110 FC scores from the two conditions, with genes sorted by their genePC1 loadings. Strongly covarying gene clusters appear at both ends of the loading spectrum, and the two ends are anti-correlated. (D) Mean ΔFC across the top 500 (yellow) and bottom 500 (green) genes. ΔFC was positive for the top genes (0.167; one-sample $t(54) = 2.07$, $P = 0.043$) and negative for the bottom genes (−0.137; $t(54) = -2.05$, $P = 0.045$), and the two sets differed (paired $t(54) = 2.08$, $P = 0.042$). (E) Mean FC across the top 500 genes. These genes were up-regulated following OT (mean FC = 0.151; one-sample $t(54) = 2.42$, $P = 0.019$) but did not change after OC (−0.016; $t(54) = -0.30$, $P = 0.76$), and OT was greater than OC within participants (paired $t(54) = 2.07$, $P = 0.043$). (F) Mean FC across the bottom 500 genes. These genes were down-regulated following OT (mean FC = −0.118; $t(54) = -2.30$, $P = 0.025$) and did not change after OC (mean FC = 0.019; $t(54) = 0.45$, $P = 0.65$), and OT was lower than OC (paired $t(54) = -2.05$, $P = 0.045$). Bars show mean ± SEM, and dots and lines show individual participants. *$P < 0.05$; *n.s.*, not significant.

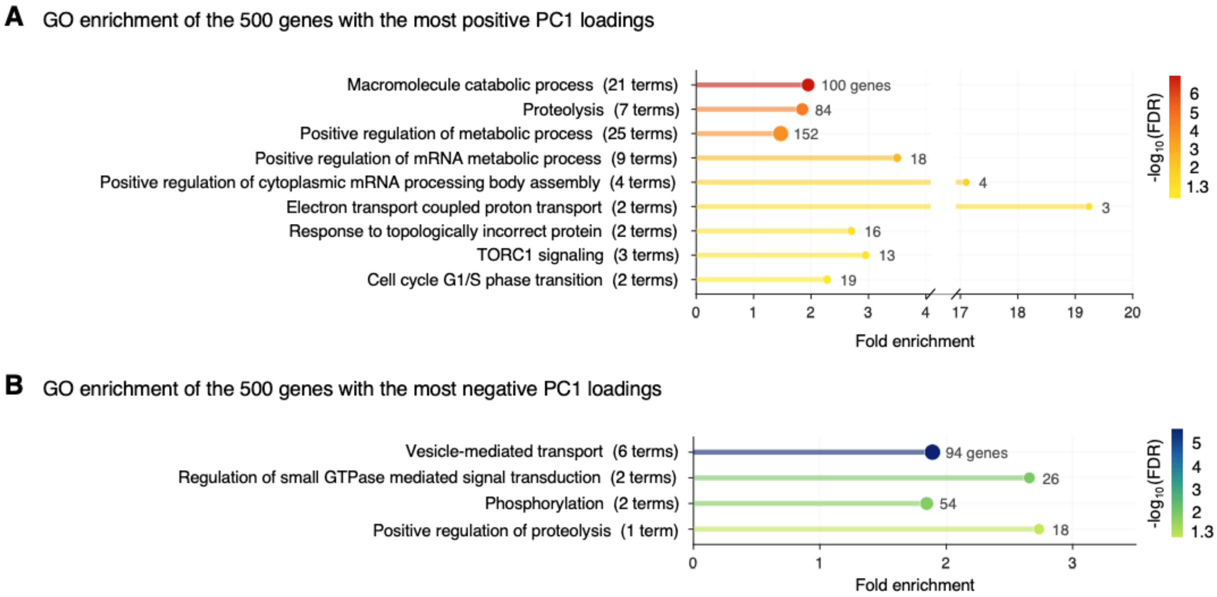


**Figure S4. Gene ontology enrichment of the genes with the strongest genePC1 loadings.** Gene ontology (GO) biological process terms enriched among (A) the top 500 genes with the most positive genePC1 loadings and (B) the bottom 500 genes with the most negative loadings (**Figure S3**), tested with ShinyGO against all 17,229 expressed genes (FDR q < 0.05, ≥ 3 genes per term). Significant terms were grouped by semantic similarity (rrvgo; Rel measure, threshold 0.7), and each bar shows the representative term of one group, with the number of grouped terms in parentheses. Bar length, fold enrichment; color, -log10(FDR); dot size and number, genes in the representative term. (A) The top 500 genes, which include 9 of the 12 mtDNA-encoded mRNAs (except *MT-ND6*), were enriched for macromolecule catabolism, proteolysis and metabolic regulation (77 terms in 9 groups). The group with the highest fold enrichment, electron transport coupled proton transport (fold enrichment = 19.2, FDR $q$ = 0.017), consists entirely of mtDNA-encoded genes (*MT-ND4*, *MT-ND5*, and *MT-CYB*), placing the mitochondrial respiratory chain at the positive end of genePC1. (B) The bottom 500 genes were enriched for vesicle-mediated transport, regulation of small GTPase-mediated signal transduction, phosphorylation and positive regulation of proteolysis (11 terms in 4 groups), and none of these terms included mtDNA-encoded genes.

| | Condition | | | | Baseline (pre) | | | | Condition × Baseline | | | | |
|---|---|---|---|---|---|---|---|---|---|---|---|---|---|
| Transcript | β | SE | *P* | *q* | β | SE | *P* | *q* | β | SE | *df* | *P* | *q* |
| MT-ATP6 | -0.224 | 0.124 | 0.073 | 0.110 | 0.393 | 0.091 | < .001 *** | < .001 | 0.252 | 0.090 | 103.0 | 0.006 ** | 0.010 |
| MT-ATP8 | -0.227 | 0.125 | 0.075 | 0.110 | 0.408 | 0.095 | < .001 *** | < .001 | 0.257 | 0.094 | 101.7 | 0.007 ** | 0.010 |
| MT-CO1 | -0.155 | 0.087 | 0.076 | 0.110 | 0.444 | 0.089 | < .001 *** | < .001 | 0.251 | 0.088 | 103.0 | 0.005 ** | 0.010 |
| MT-CO2 | -0.139 | 0.072 | 0.058 | 0.110 | 0.472 | 0.085 | < .001 *** | < .001 | 0.216 | 0.084 | 103.0 | 0.012 * | 0.014 |
| MT-CO3 | -0.152 | 0.083 | 0.071 | 0.110 | 0.434 | 0.088 | < .001 *** | < .001 | 0.239 | 0.087 | 103.0 | 0.007 ** | 0.010 |
| MT-CYB | -0.170 | 0.094 | 0.072 | 0.110 | 0.447 | 0.091 | < .001 *** | < .001 | 0.265 | 0.090 | 103.0 | 0.004 ** | 0.010 |
| MT-ND1 | -0.141 | 0.090 | 0.122 | 0.132 | 0.443 | 0.086 | < .001 *** | < .001 | 0.211 | 0.086 | 103.0 | 0.016 * | 0.017 |
| MT-ND2 | -0.199 | 0.110 | 0.072 | 0.110 | 0.444 | 0.088 | < .001 *** | < .001 | 0.247 | 0.087 | 103.0 | 0.005 ** | 0.010 |
| MT-ND3 | -0.189 | 0.112 | 0.094 | 0.122 | 0.447 | 0.087 | < .001 *** | < .001 | 0.269 | 0.087 | 103.0 | 0.002 ** | 0.010 |
| MT-ND4 | -0.188 | 0.105 | 0.076 | 0.110 | 0.424 | 0.091 | < .001 *** | < .001 | 0.255 | 0.090 | 103.0 | 0.006 ** | 0.010 |
| MT-ND4L | -0.162 | 0.102 | 0.116 | 0.132 | 0.441 | 0.090 | < .001 *** | < .001 | 0.246 | 0.090 | 103.0 | 0.007 ** | 0.010 |
| MT-ND5 | -0.183 | 0.099 | 0.068 | 0.110 | 0.435 | 0.091 | < .001 *** | < .001 | 0.276 | 0.091 | 103.0 | 0.003 ** | 0.010 |
| MT-ND6 | -0.018 | 0.068 | 0.795 | 0.795 | 0.073 | 0.094 | 0.439 | 0.439 | 0.052 | 0.097 | 102.9 | 0.589 | 0.589 |
| MT Composite | -0.177 | 0.099 | 0.077 | — | 0.431 | 0.090 | < .001 *** | — | 0.254 | 0.089 | 103.0 | 0.005 ** | — |

**Table S1. Linear mixed-effects (LME) model results for mtDNA-encoded transcripts and the MT composite.** LME model: post ~ condition × pre + order + sex + age + (1 | participant) ($n$ = 55). Condition was coded OT = −1, OC = +1 and baseline expression was mean-centered. The MT composite comprises the 12 heavy-strand protein-coding transcripts (excluding MT-ND6). Models were fitted using restricted maximum likelihood with Satterthwaite degrees of freedom. Benjamini-Hochberg (BH) FDR correction ($q$) was applied across the 13 individual transcripts. Significance markers indicate $q$ values for individual transcripts and unadjusted $P$ values for the MT composite. All mt-mRNAs except MT-ND6 showed significant baseline main effects and condition × baseline interactions after FDR correction ($q < 0.05$). Although no condition main effects survived FDR correction, their consistently negative estimates indicated a tendency toward higher post-viewing expression under OT than OC at the mean baseline. *$q < 0.05$. **$q < 0.01$. ***$q < 0.001$.

| MitoPathway | Condition | | | | Baseline (pre) | | | | Condition × Baseline | | | | |
|---|---|---|---|---|---|---|---|---|---|---|---|---|---|
| | β | SE | *P* | *q* | β | SE | *P* | *q* | β | SE | *df* | *P* | *q* |
| **Metabolism** | | | | | | | | | | | | | |
| Metabolism | 0.013 | 0.005 | 0.010 | 0.089 | 0.439 | 0.081 | < .001 *** | < .001 | 0.247 | 0.080 | 102.8 | 0.003 | 0.057 |
| Carbohydrate metabolism | 0.012 | 0.006 | 0.042 | 0.219 | 0.449 | 0.066 | < .001 *** | < .001 | 0.150 | 0.066 | 103.0 | 0.025 | 0.103 |
| Gluconeogenesis | 0.040 | 0.015 | 0.011 | 0.089 | 0.329 | 0.085 | < .001 *** | < .001 | 0.200 | 0.081 | 89.4 | 0.016 | 0.087 |
| Malate-aspartate shuttle | 0.017 | 0.011 | 0.121 | 0.321 | 0.382 | 0.079 | < .001 *** | < .001 | 0.206 | 0.080 | 103.0 | 0.011 | 0.087 |
| Glycerol phosphate shuttle | -0.013 | 0.022 | 0.574 | 0.791 | 0.527 | 0.070 | < .001 *** | < .001 | 0.060 | 0.066 | 82.7 | 0.371 | 0.564 |
| Pyruvate metabolism | -0.001 | 0.013 | 0.950 | 0.988 | 0.299 | 0.087 | < .001 *** | < .001 | 0.154 | 0.085 | 103.0 | 0.072 | 0.193 |
| TCA cycle | 0.017 | 0.010 | 0.095 | 0.305 | 0.451 | 0.071 | < .001 *** | < .001 | 0.217 | 0.072 | 101.8 | 0.003 | 0.058 |
| TCA-associated | 0.026 | 0.010 | 0.010 | 0.089 | 0.269 | 0.101 | 0.009 ** | 0.009 | 0.073 | 0.090 | 75.0 | 0.421 | 0.620 |
| Itaconate metabolism | 0.002 | 0.013 | 0.888 | 0.952 | 0.330 | 0.095 | < .001 *** | < .001 | -0.094 | 0.094 | 96.9 | 0.317 | 0.531 |
| Ketone metabolism | 0.012 | 0.013 | 0.352 | 0.576 | 0.501 | 0.057 | < .001 *** | < .001 | 0.112 | 0.056 | 99.1 | 0.048 | 0.139 |
| Propanoate metabolism | -0.007 | 0.016 | 0.653 | 0.856 | 0.355 | 0.072 | < .001 *** | < .001 | -0.022 | 0.071 | 103.0 | 0.754 | 0.871 |
| Lipid metabolism | 0.008 | 0.005 | 0.121 | 0.321 | 0.317 | 0.073 | < .001 *** | < .001 | 0.203 | 0.072 | 94.3 | 0.006 | 0.074 |
| Fatty acid oxidation | 0.009 | 0.008 | 0.248 | 0.465 | 0.298 | 0.077 | < .001 *** | < .001 | 0.210 | 0.078 | 98.4 | 0.008 | 0.087 |
| Type II fatty acid synthesis | -0.008 | 0.012 | 0.494 | 0.698 | 0.379 | 0.062 | < .001 *** | < .001 | 0.072 | 0.060 | 99.8 | 0.238 | 0.439 |
| Lipoate insertion | -0.031 | 0.027 | 0.247 | 0.465 | 0.397 | 0.087 | < .001 *** | < .001 | 0.194 | 0.086 | 95.4 | 0.026 | 0.103 |
| Cardiolipin synthesis | 0.011 | 0.012 | 0.344 | 0.569 | 0.295 | 0.061 | < .001 *** | < .001 | 0.055 | 0.061 | 103.0 | 0.371 | 0.564 |
| Cholesterol, bile acid, steroid synthesis | 0.039 | 0.016 | 0.018 | 0.131 | 0.319 | 0.087 | < .001 *** | < .001 | 0.219 | 0.084 | 91.8 | 0.011 | 0.087 |
| Cholesterol-associated | 0.006 | 0.014 | 0.676 | 0.857 | 0.304 | 0.095 | 0.002 ** | 0.002 | 0.094 | 0.096 | 103.0 | 0.326 | 0.540 |
| Phospholipid metabolism | 0.002 | 0.008 | 0.817 | 0.939 | 0.366 | 0.061 | < .001 *** | < .001 | 0.107 | 0.060 | 89.2 | 0.080 | 0.205 |
| Eicosanoid metabolism | 0.026 | 0.013 | 0.042 | 0.219 | 0.393 | 0.094 | < .001 *** | < .001 | 0.070 | 0.092 | 93.6 | 0.450 | 0.653 |
| Amino acid metabolism | 0.018 | 0.008 | 0.024 | 0.167 | 0.550 | 0.080 | < .001 *** | < .001 | 0.077 | 0.080 | 103.0 | 0.336 | 0.545 |
| Branched-chain amino acid metabolism | 0.009 | 0.008 | 0.284 | 0.503 | 0.436 | 0.072 | < .001 *** | < .001 | 0.099 | 0.072 | 103.0 | 0.175 | 0.356 |
| Branched-chain amino acid dehydrogenase complex | -0.005 | 0.012 | 0.694 | 0.862 | 0.223 | 0.086 | 0.011 * | 0.011 | -0.045 | 0.086 | 103.0 | 0.599 | 0.790 |
| Lysine metabolism | 0.016 | 0.008 | 0.047 | 0.219 | 0.432 | 0.078 | < .001 *** | < .001 | 0.041 | 0.079 | 101.3 | 0.607 | 0.793 |
| Serine metabolism | 0.046 | 0.020 | 0.030 | 0.197 | 0.340 | 0.108 | 0.002 ** | 0.003 | 0.150 | 0.107 | 96.9 | 0.162 | 0.344 |
| Glycine metabolism | 0.033 | 0.019 | 0.080 | 0.277 | 0.637 | 0.077 | < .001 *** | < .001 | -0.048 | 0.077 | 103.0 | 0.538 | 0.716 |
| Glycine cleavage system | 0.000 | 0.043 | 0.999 | 0.999 | 0.633 | 0.070 | < .001 *** | < .001 | 0.028 | 0.070 | 103.0 | 0.685 | 0.823 |
| Glutamate metabolism | 0.019 | 0.018 | 0.289 | 0.506 | 0.251 | 0.083 | 0.003 ** | 0.004 | 0.007 | 0.083 | 102.4 | 0.937 | 0.984 |
| Proline metabolism | 0.054 | 0.027 | 0.050 | 0.225 | 0.800 | 0.081 | < .001 *** | < .001 | 0.107 | 0.075 | 85.5 | 0.159 | 0.344 |
| Glyoxylate metabolism | 0.026 | 0.017 | 0.141 | 0.350 | 0.327 | 0.092 | < .001 *** | < .001 | 0.140 | 0.092 | 96.9 | 0.129 | 0.296 |
| GABA metabolism | 0.019 | 0.012 | 0.112 | 0.316 | 0.343 | 0.077 | < .001 *** | < .001 | 0.101 | 0.073 | 86.6 | 0.166 | 0.344 |
| Catechol metabolism | 0.059 | 0.031 | 0.066 | 0.246 | 0.467 | 0.090 | < .001 *** | < .001 | 0.181 | 0.089 | 102.3 | 0.045 | 0.137 |
| Kynurenine metabolism | -0.003 | 0.017 | 0.852 | 0.939 | 0.369 | 0.081 | < .001 *** | < .001 | 0.175 | 0.072 | 75.6 | 0.017 | 0.088 |
| Urea cycle | -0.005 | 0.013 | 0.676 | 0.857 | 0.327 | 0.077 | < .001 *** | < .001 | -0.033 | 0.076 | 98.3 | 0.670 | 0.819 |
| Nucleotide metabolism | 0.041 | 0.010 | < .001 ** | 0.008 | 0.543 | 0.097 | < .001 *** | < .001 | 0.196 | 0.096 | 97.4 | 0.045 | 0.137 |
| Nucleotide import | 0.067 | 0.018 | < .001 * | 0.017 | 0.713 | 0.083 | < .001 *** | < .001 | -0.087 | 0.079 | 95.3 | 0.272 | 0.480 |
| Nucleotide synthesis and processing | 0.033 | 0.010 | 0.001 * | 0.024 | 0.541 | 0.104 | < .001 *** | < .001 | 0.166 | 0.104 | 96.4 | 0.113 | 0.268 |
| Creatine metabolism | 0.036 | 0.048 | 0.455 | 0.659 | 0.521 | 0.059 | < .001 *** | < .001 | 0.079 | 0.048 | 54.8 | 0.103 | 0.253 |
| Metals and cofactors | 0.006 | 0.005 | 0.253 | 0.465 | 0.334 | 0.080 | < .001 *** | < .001 | 0.171 | 0.079 | 103.0 | 0.033 | 0.115 |
| Carnitine synthesis and transport | 0.023 | 0.014 | 0.105 | 0.312 | 0.276 | 0.081 | < .001 ** | 0.001 | 0.048 | 0.076 | 79.1 | 0.528 | 0.716 |
| Carnitine shuttle | 0.029 | 0.017 | 0.095 | 0.305 | 0.253 | 0.090 | 0.006 ** | 0.007 | 0.038 | 0.083 | 78.4 | 0.650 | 0.814 |
| Coenzyme A metabolism | 0.038 | 0.013 | 0.007 | 0.085 | 0.431 | 0.090 | < .001 *** | < .001 | -0.060 | 0.081 | 76.7 | 0.465 | 0.666 |
| Coenzyme Q metabolism | 0.014 | 0.008 | 0.066 | 0.246 | 0.345 | 0.070 | < .001 *** | < .001 | 0.046 | 0.069 | 102.7 | 0.504 | 0.701 |
| Copper metabolism | -0.012 | 0.010 | 0.218 | 0.444 | 0.577 | 0.064 | < .001 *** | < .001 | 0.032 | 0.065 | 103.0 | 0.617 | 0.793 |
| Iron homeostasis | 0.005 | 0.010 | 0.583 | 0.796 | 0.668 | 0.057 | < .001 *** | < .001 | 0.047 | 0.056 | 91.4 | 0.399 | 0.601 |
| Heme synthesis and processing | -0.002 | 0.008 | 0.777 | 0.918 | 0.257 | 0.072 | < .001 *** | < .001 | 0.007 | 0.073 | 100.3 | 0.924 | 0.983 |
| Fe-S cluster biosynthesis | 0.004 | 0.011 | 0.710 | 0.867 | 0.401 | 0.087 | < .001 *** | < .001 | 0.163 | 0.084 | 103.0 | 0.055 | 0.155 |
| Molybdenum cofactor synthesis and proteins | -0.007 | 0.019 | 0.717 | 0.868 | 0.200 | 0.077 | 0.011 * | 0.011 | -0.004 | 0.074 | 78.5 | 0.953 | 0.993 |
| NAD biosynthesis and metabolism | -0.006 | 0.007 | 0.388 | 0.609 | 0.260 | 0.069 | < .001 *** | < .001 | 0.166 | 0.066 | 88.7 | 0.014 | 0.087 |
| Tetrahydrobiopterin synthesis | 0.010 | 0.023 | 0.655 | 0.856 | 0.261 | 0.092 | 0.006 ** | 0.006 | 0.120 | 0.086 | 83.3 | 0.165 | 0.344 |
| Fe-S-containing proteins | 0.006 | 0.008 | 0.495 | 0.698 | 0.419 | 0.088 | < .001 *** | < .001 | 0.287 | 0.087 | 103.0 | 0.001 * | 0.048 |
| Heme-containing proteins | 0.027 | 0.016 | 0.107 | 0.312 | 0.361 | 0.084 | < .001 *** | < .001 | -0.079 | 0.082 | 90.3 | 0.336 | 0.545 |
| Vitamin metabolism | 0.007 | 0.007 | 0.371 | 0.595 | 0.401 | 0.078 | < .001 *** | < .001 | 0.007 | 0.076 | 102.6 | 0.930 | 0.983 |
| Choline and betaine metabolism | 0.036 | 0.021 | 0.096 | 0.305 | 0.306 | 0.091 | 0.001 ** | 0.001 | 0.056 | 0.091 | 102.8 | 0.536 | 0.716 |
| Biotin utilizing proteins | 0.006 | 0.014 | 0.679 | 0.857 | 0.593 | 0.072 | < .001 *** | < .001 | 0.196 | 0.072 | 97.5 | 0.008 | 0.087 |
| Vitamin A metabolism | -0.034 | 0.013 | 0.013 | 0.105 | 0.293 | 0.069 | < .001 *** | < .001 | 0.031 | 0.067 | 82.6 | 0.649 | 0.814 |
| Vitamin B1 metabolism | -0.002 | 0.026 | 0.933 | 0.986 | 0.249 | 0.092 | 0.008 ** | 0.009 | 0.130 | 0.090 | 103.0 | 0.150 | 0.339 |
| Vitamin B2 metabolism | 0.000 | 0.021 | 0.993 | 0.999 | 0.119 | 0.069 | 0.086 | 0.088 | 0.025 | 0.069 | 103.0 | 0.719 | 0.857 |
| Vitamin B6 metabolism | 0.026 | 0.015 | 0.100 | 0.311 | 0.301 | 0.081 | < .001 *** | < .001 | 0.189 | 0.082 | 101.5 | 0.023 | 0.099 |
| Vitamin B12 metabolism | 0.017 | 0.016 | 0.275 | 0.495 | 0.418 | 0.083 | < .001 *** | < .001 | -0.075 | 0.082 | 103.0 | 0.360 | 0.564 |
| Vitamin C metabolism | -0.020 | 0.013 | 0.137 | 0.346 | 0.317 | 0.106 | 0.004 ** | 0.004 | 0.161 | 0.096 | 72.8 | 0.097 | 0.242 |
| Vitamin D metabolism | 0.048 | 0.030 | 0.119 | 0.321 | 0.112 | 0.091 | 0.222 | 0.222 | -0.112 | 0.091 | 92.1 | 0.221 | 0.423 |
| Folate and 1-C metabolism | 0.004 | 0.012 | 0.758 | 0.910 | 0.474 | 0.073 | < .001 *** | < .001 | -0.014 | 0.071 | 100.6 | 0.847 | 0.928 |
| Detoxification | 0.035 | 0.010 | 0.001 * | 0.024 | 0.481 | 0.092 | < .001 *** | < .001 | 0.213 | 0.092 | 97.6 | 0.023 | 0.099 |
| Xenobiotic metabolism | 0.035 | 0.013 | 0.008 | 0.085 | 0.454 | 0.092 | < .001 *** | < .001 | 0.112 | 0.090 | 92.7 | 0.219 | 0.423 |
| ROS and glutathione metabolism | 0.037 | 0.011 | 0.002 * | 0.028 | 0.493 | 0.083 | < .001 *** | < .001 | 0.218 | 0.084 | 97.6 | 0.011 | 0.087 |
| Amidoxime reducing complex | 0.004 | 0.020 | 0.843 | 0.939 | 0.166 | 0.079 | 0.038 * | 0.040 | 0.036 | 0.079 | 85.9 | 0.647 | 0.814 |
| Selenoproteins | 0.060 | 0.022 | 0.009 | 0.085 | 0.482 | 0.093 | < .001 *** | < .001 | 0.159 | 0.090 | 99.7 | 0.080 | 0.205 |

| | | | | | | | | | | | | | |
|---|---|---|---|---|---|---|---|---|---|---|---|---|---|
| Electron carriers | 0.002 | 0.011 | 0.848 | 0.939 | 0.421 | 0.080 | < .001 *** | < .001 | 0.023 | 0.077 | 94.1 | 0.769 | 0.874 |
| Cytochromes | 0.013 | 0.015 | 0.416 | 0.622 | 0.476 | 0.082 | < .001 *** | < .001 | 0.090 | 0.080 | 90.4 | 0.268 | 0.480 |
| Q-linked reactions, other | -0.002 | 0.011 | 0.885 | 0.952 | 0.340 | 0.077 | < .001 *** | < .001 | -0.001 | 0.074 | 90.7 | 0.995 | 1.000 |
| Sulfur metabolism | 0.025 | 0.013 | 0.055 | 0.230 | 0.172 | 0.088 | 0.055 | 0.056 | 0.116 | 0.087 | 95.2 | 0.188 | 0.379 |
| **OXPHOS** | | | | | | | | | | | | | |
| OXPHOS | 0.011 | 0.009 | 0.223 | 0.444 | 0.439 | 0.082 | < .001 *** | < .001 | 0.191 | 0.084 | 102.8 | 0.025 | 0.103 |
| OXPHOS subunits | 0.024 | 0.012 | 0.056 | 0.230 | 0.461 | 0.083 | < .001 *** | < .001 | 0.176 | 0.084 | 99.9 | 0.039 | 0.130 |
| OXPHOS assembly factors | -0.002 | 0.007 | 0.783 | 0.918 | 0.338 | 0.078 | < .001 *** | < .001 | 0.228 | 0.078 | 103.0 | 0.004 | 0.070 |
| Complex I | 0.014 | 0.009 | 0.131 | 0.343 | 0.447 | 0.093 | < .001 *** | < .001 | 0.213 | 0.095 | 103.0 | 0.027 | 0.103 |
| CI subunits | 0.019 | 0.010 | 0.064 | 0.246 | 0.455 | 0.097 | < .001 *** | < .001 | 0.135 | 0.097 | 96.8 | 0.166 | 0.344 |
| CI assembly factors | 0.007 | 0.009 | 0.443 | 0.646 | 0.335 | 0.087 | < .001 *** | < .001 | 0.210 | 0.087 | 103.0 | 0.018 | 0.088 |
| Complex II | 0.010 | 0.011 | 0.323 | 0.547 | 0.262 | 0.087 | 0.004 ** | 0.004 | 0.149 | 0.089 | 95.3 | 0.097 | 0.242 |
| CII subunits | 0.007 | 0.013 | 0.598 | 0.808 | 0.331 | 0.083 | < .001 *** | < .001 | 0.076 | 0.081 | 96.2 | 0.351 | 0.562 |
| CII assembly factors | 0.011 | 0.016 | 0.497 | 0.698 | 0.158 | 0.106 | 0.141 | 0.142 | 0.091 | 0.108 | 101.1 | 0.406 | 0.605 |
| Complex III | 0.024 | 0.011 | 0.036 | 0.219 | 0.473 | 0.103 | < .001 *** | < .001 | 0.189 | 0.104 | 102.3 | 0.073 | 0.193 |
| CIII subunits | 0.043 | 0.016 | 0.008 | 0.085 | 0.488 | 0.086 | < .001 *** | < .001 | 0.217 | 0.087 | 96.7 | 0.014 | 0.087 |
| CIII assembly factors | -0.004 | 0.012 | 0.707 | 0.867 | 0.367 | 0.095 | < .001 *** | < .001 | 0.061 | 0.088 | 94.4 | 0.493 | 0.699 |
| Complex IV | 0.002 | 0.009 | 0.837 | 0.939 | 0.443 | 0.075 | < .001 *** | < .001 | 0.167 | 0.075 | 100.7 | 0.029 | 0.103 |
| CIV subunits | 0.035 | 0.019 | 0.068 | 0.247 | 0.391 | 0.077 | < .001 *** | < .001 | 0.194 | 0.078 | 99.9 | 0.015 | 0.087 |
| CIV assembly factors | -0.011 | 0.008 | 0.152 | 0.357 | 0.440 | 0.077 | < .001 *** | < .001 | 0.172 | 0.076 | 95.9 | 0.027 | 0.103 |
| Complex V | 0.013 | 0.013 | 0.315 | 0.539 | 0.370 | 0.076 | < .001 *** | < .001 | 0.163 | 0.078 | 103.0 | 0.039 | 0.130 |
| CV subunits | 0.026 | 0.015 | 0.094 | 0.305 | 0.371 | 0.081 | < .001 *** | < .001 | 0.161 | 0.083 | 102.7 | 0.054 | 0.155 |
| CV assembly factors | -0.024 | 0.017 | 0.154 | 0.357 | 0.220 | 0.076 | 0.005 ** | 0.005 | 0.161 | 0.078 | 101.0 | 0.041 | 0.132 |
| Respirasome assembly | 0.029 | 0.018 | 0.109 | 0.312 | 0.387 | 0.079 | < .001 *** | < .001 | 0.180 | 0.076 | 93.2 | 0.021 | 0.096 |
| Cytochrome C | -0.005 | 0.030 | 0.863 | 0.939 | 0.345 | 0.089 | < .001 *** | < .001 | 0.083 | 0.089 | 103.0 | 0.356 | 0.564 |
| **Mitochondrial central dogma** | | | | | | | | | | | | | |
| Mitochondrial central dogma | 0.002 | 0.007 | 0.763 | 0.910 | 0.532 | 0.081 | < .001 *** | < .001 | 0.227 | 0.079 | 103.0 | 0.005 | 0.074 |
| mtDNA maintenance | 0.010 | 0.008 | 0.219 | 0.444 | 0.606 | 0.066 | < .001 *** | < .001 | 0.001 | 0.060 | 89.8 | 0.986 | 1.000 |
| mtDNA replication | 0.000 | 0.006 | 0.977 | 0.997 | 0.361 | 0.076 | < .001 *** | < .001 | 0.169 | 0.071 | 92.1 | 0.019 | 0.094 |
| mtDNA nucleoid | 0.017 | 0.008 | 0.046 | 0.219 | 0.402 | 0.078 | < .001 *** | < .001 | 0.024 | 0.073 | 81.4 | 0.745 | 0.871 |
| mtDNA repair | 0.008 | 0.009 | 0.385 | 0.609 | 0.521 | 0.071 | < .001 *** | < .001 | 0.082 | 0.063 | 74.3 | 0.196 | 0.390 |
| mtDNA modifications | -0.037 | 0.032 | 0.242 | 0.465 | 0.262 | 0.087 | 0.003 ** | 0.004 | 0.170 | 0.085 | 103.0 | 0.047 | 0.139 |
| mtDNA stability and decay | 0.003 | 0.014 | 0.827 | 0.939 | 0.189 | 0.109 | 0.085 | 0.087 | 0.074 | 0.109 | 100.5 | 0.498 | 0.699 |
| mtRNA metabolism | -0.006 | 0.006 | 0.301 | 0.522 | 0.452 | 0.079 | < .001 *** | < .001 | 0.245 | 0.077 | 103.0 | 0.002 | 0.051 |
| Transcription | 0.008 | 0.008 | 0.333 | 0.557 | 0.501 | 0.086 | < .001 *** | < .001 | -0.022 | 0.081 | 84.7 | 0.786 | 0.874 |
| mtRNA granules | 0.000 | 0.007 | 0.960 | 0.988 | 0.375 | 0.093 | < .001 *** | < .001 | 0.223 | 0.091 | 103.0 | 0.016 | 0.087 |
| Polycistronic mtRNA processing | -0.010 | 0.009 | 0.250 | 0.465 | 0.558 | 0.085 | < .001 *** | < .001 | 0.076 | 0.084 | 103.0 | 0.366 | 0.564 |
| mt-tRNA modifications | -0.006 | 0.007 | 0.417 | 0.622 | 0.345 | 0.085 | < .001 *** | < .001 | 0.136 | 0.085 | 103.0 | 0.112 | 0.268 |
| mt-rRNA modifications | -0.001 | 0.007 | 0.854 | 0.939 | 0.207 | 0.066 | 0.002 ** | 0.003 | 0.160 | 0.065 | 98.8 | 0.015 | 0.087 |
| mt-mRNA modifications | -0.016 | 0.013 | 0.224 | 0.444 | 0.183 | 0.088 | 0.041 * | 0.043 | 0.192 | 0.086 | 94.0 | 0.028 | 0.103 |
| mtRNA stability and decay | -0.010 | 0.013 | 0.414 | 0.622 | 0.348 | 0.089 | < .001 *** | < .001 | 0.179 | 0.087 | 103.0 | 0.044 | 0.137 |
| Translation | 0.004 | 0.008 | 0.625 | 0.832 | 0.491 | 0.089 | < .001 *** | < .001 | 0.227 | 0.089 | 103.0 | 0.012 | 0.087 |
| Mitochondrial ribosome | 0.014 | 0.011 | 0.241 | 0.465 | 0.416 | 0.091 | < .001 *** | < .001 | 0.144 | 0.092 | 103.0 | 0.122 | 0.283 |
| Mitochondrial ribosome assembly | -0.013 | 0.006 | 0.053 | 0.230 | 0.295 | 0.066 | < .001 *** | < .001 | 0.157 | 0.064 | 102.2 | 0.016 | 0.087 |
| Translation factors | -0.012 | 0.010 | 0.222 | 0.444 | 0.375 | 0.081 | < .001 *** | < .001 | 0.214 | 0.082 | 103.0 | 0.010 | 0.087 |
| mt-tRNA synthetases | 0.005 | 0.008 | 0.533 | 0.742 | 0.562 | 0.086 | < .001 *** | < .001 | 0.303 | 0.081 | 99.8 | < .001 * | 0.015 |
| fMet processing | -0.005 | 0.026 | 0.852 | 0.939 | 0.278 | 0.083 | 0.001 ** | 0.001 | 0.080 | 0.079 | 103.0 | 0.311 | 0.526 |
| **Small molecule transport** | | | | | | | | | | | | | |
| Small molecule transport | 0.011 | 0.006 | 0.061 | 0.244 | 0.409 | 0.079 | < .001 *** | < .001 | 0.002 | 0.073 | 94.4 | 0.981 | 1.000 |
| SLC25A family | 0.012 | 0.006 | 0.043 | 0.219 | 0.437 | 0.080 | < .001 *** | < .001 | 0.000 | 0.075 | 88.7 | 1.000 | 1.000 |
| ABC transporters | 0.003 | 0.016 | 0.860 | 0.939 | 0.340 | 0.084 | < .001 *** | < .001 | 0.090 | 0.079 | 91.9 | 0.258 | 0.469 |
| Sideroflexins | -0.004 | 0.009 | 0.690 | 0.862 | 0.484 | 0.074 | < .001 *** | < .001 | 0.074 | 0.067 | 82.6 | 0.274 | 0.480 |
| Calcium uniporter | 0.023 | 0.026 | 0.371 | 0.595 | 0.286 | 0.123 | 0.022 * | 0.023 | -0.054 | 0.124 | 103.0 | 0.665 | 0.819 |
| **Mitochondrial dynamics and surveillance** | | | | | | | | | | | | | |
| Mitochondrial dynamics and surveillance | 0.018 | 0.006 | 0.004 | 0.061 | 0.440 | 0.081 | < .001 *** | < .001 | 0.085 | 0.081 | 96.4 | 0.298 | 0.516 |
| Fusion | -0.001 | 0.012 | 0.958 | 0.988 | 0.256 | 0.090 | 0.006 ** | 0.006 | 0.022 | 0.090 | 99.3 | 0.806 | 0.890 |
| Fission | 0.007 | 0.008 | 0.403 | 0.622 | 0.354 | 0.077 | < .001 *** | < .001 | -0.001 | 0.075 | 87.0 | 0.987 | 1.000 |
| Organelle contact sites | 0.032 | 0.008 | < .001 ** | 0.008 | 0.555 | 0.105 | < .001 *** | < .001 | 0.320 | 0.100 | 88.4 | 0.002 | 0.051 |
| Intramitochondrial membrane interactions | 0.004 | 0.009 | 0.677 | 0.857 | 0.587 | 0.070 | < .001 *** | < .001 | 0.008 | 0.070 | 103.0 | 0.907 | 0.972 |
| Trafficking | 0.000 | 0.008 | 0.962 | 0.988 | 0.365 | 0.075 | < .001 *** | < .001 | -0.056 | 0.075 | 97.9 | 0.451 | 0.653 |
| Mitophagy | 0.021 | 0.010 | 0.042 | 0.219 | 0.402 | 0.068 | < .001 *** | < .001 | 0.318 | 0.072 | 101.5 | < .001 ** | 0.002 |
| Autophagy | 0.015 | 0.007 | 0.047 | 0.219 | 0.437 | 0.065 | < .001 *** | < .001 | 0.306 | 0.067 | 94.7 | < .001 ** | 0.002 |
| Apoptosis | 0.027 | 0.010 | 0.007 | 0.085 | 0.334 | 0.084 | < .001 *** | < .001 | 0.047 | 0.075 | 75.1 | 0.536 | 0.716 |
| Cristae formation | 0.016 | 0.014 | 0.259 | 0.470 | 0.333 | 0.081 | < .001 *** | < .001 | -0.041 | 0.080 | 103.0 | 0.612 | 0.793 |
| MICOS complex | 0.032 | 0.015 | 0.040 | 0.219 | 0.365 | 0.081 | < .001 *** | < .001 | -0.095 | 0.080 | 103.0 | 0.238 | 0.439 |
| **Protein import, sorting and homeostasis** | | | | | | | | | | | | | |
| Protein import, sorting and homeostasis | 0.010 | 0.007 | 0.146 | 0.357 | 0.418 | 0.077 | < .001 *** | < .001 | 0.109 | 0.077 | 102.3 | 0.161 | 0.344 |
| Protein import and sorting | 0.009 | 0.007 | 0.200 | 0.444 | 0.316 | 0.084 | < .001 *** | < .001 | 0.054 | 0.086 | 102.4 | 0.534 | 0.716 |
| TOM | 0.013 | 0.011 | 0.219 | 0.444 | 0.273 | 0.077 | < .001 *** | < .001 | 0.181 | 0.082 | 101.6 | 0.029 | 0.103 |
| SAM | 0.007 | 0.013 | 0.602 | 0.808 | 0.370 | 0.086 | < .001 *** | < .001 | 0.024 | 0.086 | 101.9 | 0.786 | 0.874 |
| MIA40 | 0.025 | 0.019 | 0.194 | 0.439 | 0.256 | 0.088 | 0.004 ** | 0.005 | 0.249 | 0.089 | 100.9 | 0.006 | 0.074 |

| | | | | | | | | | | | | | | |
|---|---|---|---|---|---|---|---|---|---|---|---|---|---|---|
| TIM22 carrier pathway | 0.023 | 0.015 | 0.136 | | 0.346 | 0.382 | 0.106 | < .001 | *** | < .001 | -0.032 | 0.102 | 88.6 | 0.754 |
| TIM23 presequence pathway | 0.022 | 0.012 | 0.079 | | 0.277 | 0.346 | 0.089 | < .001 | *** | < .001 | 0.094 | 0.091 | 101.2 | 0.303 |
| Import motor | -0.012 | 0.015 | 0.427 | | 0.630 | 0.406 | 0.085 | < .001 | *** | < .001 | 0.202 | 0.083 | 103.0 | 0.016 |
| OXA | -0.013 | 0.016 | 0.406 | | 0.622 | 0.508 | 0.078 | < .001 | *** | < .001 | 0.097 | 0.078 | 97.9 | 0.217 |
| Preprotein cleavage | 0.000 | 0.013 | 0.987 | | 0.999 | 0.344 | 0.086 | < .001 | *** | < .001 | -0.014 | 0.085 | 102.9 | 0.865 |
| Protein homeostasis | 0.009 | 0.007 | 0.211 | | 0.444 | 0.414 | 0.073 | < .001 | *** | < .001 | 0.087 | 0.073 | 103.0 | 0.235 |
| Proteases | 0.001 | 0.007 | 0.906 | | 0.965 | 0.374 | 0.068 | < .001 | *** | < .001 | 0.030 | 0.066 | 103.0 | 0.656 |
| Chaperones | 0.018 | 0.011 | 0.103 | | 0.312 | 0.286 | 0.085 | 0.001 | ** | 0.001 | 0.035 | 0.086 | 103.0 | 0.685 |
| **Signaling** | | | | | | | | | | | | | | |
| Signaling | 0.033 | 0.008 | < .001 | ** | 0.008 | 0.419 | 0.098 | < .001 | *** | < .001 | 0.239 | 0.093 | 86.0 | 0.012 |
| Calcium homeostasis | 0.039 | 0.011 | < .001 | * | 0.017 | 0.519 | 0.095 | < .001 | *** | < .001 | -0.026 | 0.094 | 94.7 | 0.780 |
| Calcium cycle | 0.021 | 0.015 | 0.156 | | 0.357 | 0.273 | 0.118 | 0.022 | * | 0.023 | -0.021 | 0.117 | 102.7 | 0.858 |
| Mitochondrial permeability transition pore | 0.091 | 0.022 | < .001 | ** | 0.008 | 0.440 | 0.098 | < .001 | *** | < .001 | 0.002 | 0.095 | 96.9 | 0.987 |
| EF hand proteins | 0.020 | 0.016 | 0.214 | | 0.444 | 0.418 | 0.106 | < .001 | *** | < .001 | -0.029 | 0.104 | 93.8 | 0.780 |
| Immune response | 0.034 | 0.015 | 0.027 | | 0.185 | 0.367 | 0.088 | < .001 | *** | < .001 | 0.026 | 0.083 | 95.5 | 0.752 |
| cAMP-PKA signaling | 0.023 | 0.015 | 0.151 | | 0.357 | 0.409 | 0.082 | < .001 | *** | < .001 | 0.149 | 0.081 | 95.9 | 0.070 |

**Table S2. LME model results for all 149 mitochondrial pathways.** Pathway scores represent mean $\log_2$CPM across constituent genes (*n* = 55). Results indicate regression coefficients (β), standard errors (*SE*), *P* values, and BH FDR-adjusted *q* values for condition, baseline, and their interaction. Satterthwaite degrees of freedom (*df*) are reported. Asterisks indicate FDR-adjusted significance: *$q < 0.05$. **$q < 0.01$. ***$q < 0.001$.

| | | Main results ($n$ = 55) | | | Library size control | | Outlier controls | | | |
|---|---|---|---|---|---|---|---|---|---|---|
| Effect | MitoPathway | β | $P$ | FDR $q$ | β | $P$ | Excluded subID | $n$ | β | $P$ |
| Condition | Signaling | 0.033 | 0.0001 | 0.0077 | 0.032 | 0.0002 | 22, 53 | 53 | 0.02896 | 0.00028 |
| Condition | Mitochondrial permeability transition pore | 0.091 | 0.0002 | 0.0079 | 0.089 | 0.0002 | 12, 43, 53 | 52 | 0.0692 | 0.001 |
| Condition | Nucleotide metabolism | 0.041 | 0.0002 | 0.0079 | 0.039 | 0.0005 | | 55 | 0.04059 | 0.000 |
| Condition | Organelle contact sites | 0.032 | 0.0002 | 0.0079 | 0.031 | 0.0008 | 5, 22, 39, 40, 53, 59 | 49 | 0.01873 | 0.021 |
| Condition | Calcium homeostasis | 0.039 | 0.0006 | 0.0165 | 0.038 | 0.0009 | 53 | 54 | 0.03641 | 0.001 |
| Condition | Nucleotide import | 0.067 | 0.0007 | 0.0172 | 0.065 | 0.0011 | 53 | 54 | 0.06548 | 0.001 |
| Condition | Detoxification | 0.035 | 0.0013 | 0.0244 | 0.034 | 0.0035 | 22, 39, 48 | 52 | 0.02954 | 0.009 |
| Condition | Nucleotide synthesis and processing | 0.033 | 0.0013 | 0.0244 | 0.031 | 0.0042 | | 55 | 0.03323 | 0.001 |
| Condition | ROS and glutathione metabolism | 0.037 | 0.0017 | 0.0277 | 0.035 | 0.0037 | 22, 23 | 53 | 0.03542 | 0.002 |
| Condition × baseline | Autophagy | 0.306 | 0.0000 | 0.0019 | 0.343 | 0.0000 | 22, 48, 58, 67 | 51 | 0.29041 | 0.000 |
| Condition × baseline | Mitophagy | 0.318 | 0.0000 | 0.0019 | 0.345 | 0.0000 | 22, 58, 60, 67 | 51 | 0.30594 | 0.000 |
| Condition × baseline | mt-tRNA synthetases | 0.303 | 0.0003 | 0.0150 | 0.340 | 0.0002 | 7, 53, 67 | 52 | 0.21885 | 0.029 |
| Condition × baseline | Fe-S-containing proteins | 0.287 | 0.0013 | 0.0483 | 0.308 | 0.0007 | 7, 29, 53, 67 | 51 | 0.18382 | 0.072 |

**Table S3. Sensitivity analyses of condition-dependent mitochondrial pathway expression.** Results for pathway with significant condition main effects or condition × baseline interactions in the primary analysis ($n$ = 55). Library-size control models used normalization factors recalculated after excluding mtDNA-derived reads. Outlier-control models excluded all observations from participants with pre- or post-viewing pathway scores exceeding three scaled median absolute deviations from the condition-specific median. The same LME model was refitted for each analysis. β denotes the regression coefficients, $P$ the unadjusted significance level, and FDR $q$ the BH adjusted $P$ value from the primary analysis. Excluded subIDs column indicates removed participants; $n$ indicates the remaining sample size.

| | | | OT–OC | | OT condition | | OC condition | |
|---|---|---|---|---|---|---|---|---|
| Domain | Measure | *n* | *r* | *p* | *r* | *p* | *r* | *p* |
| Personality | Animal Attitude Scale | 55 | -0.050 | 0.719 | | | | |
| | Interpersonal Reactivity Index: Perspective Taking | 55 | -0.028 | 0.840 | | | | |
| | Interpersonal Reactivity Index: Fantasy | 55 | 0.248 | 0.068 | | | | |
| | Interpersonal Reactivity Index: Empathic Concern | 55 | 0.121 | 0.380 | | | | |
| | Interpersonal Reactivity Index: Personal Distress | 55 | -0.048 | 0.728 | | | | |
| | Interpersonal Reactivity Index: Total | 55 | 0.105 | 0.447 | | | | |
| Physiology | Heart rate during movie-viewing | 53 | 0.055 | 0.698 | -0.048 | 0.730 | -0.121 | 0.388 |
| | Skin conductance level during movie-viewing | 49 | 0.209 | 0.149 | 0.055 | 0.710 | 0.031 | 0.833 |
| Emotion ratings | Warm | 55 | 0.007 | 0.961 | -0.022 | 0.873 | 0.056 | 0.686 |
| | Joyful | 55 | 0.040 | 0.775 | -0.032 | 0.814 | 0.131 | 0.341 |
| | Inspired | 55 | -0.138 | 0.315 | -0.110 | 0.423 | 0.030 | 0.828 |
| | Ashamed | 55 | 0.050 | 0.716 | 0.019 | 0.891 | -0.090 | 0.515 |
| | Sad | 55 | 0.126 | 0.359 | 0.174 | 0.204 | 0.000 | 0.998 |
| | Horrified | 55 | 0.131 | 0.340 | 0.155 | 0.257 | 0.079 | 0.567 |
| | Disgusted | 55 | 0.186 | 0.175 | 0.165 | 0.228 | -0.027 | 0.845 |

**Table S4. Associations of MT composite expression changes with personality, autonomic responses, and emotional ratings.**

Pearson correlations between the mitochondrial (MT) composite fold change and individual-difference measures ($N = 55$). For condition-varying measures, FC was matched to the measure from the same condition. In the OT – OC columns, the within-subject difference in FC was correlated with the corresponding difference in each measure. In the OT and OC condition columns, FC and the measure were taken from that condition alone. Personality traits (Animal Attitude Scale; Interpersonal Reactivity Index) were measured once per participant, so they were correlated with ΔFC only. Heart rate (HR) and skin conductance level (SCL) were averaged over the movie-viewing runs. Participants with poor-quality physiological recordings were excluded (HR, $n = 53$; SCL, $n = 49$). Emotional ratings were averaged across the nine movie clips in each condition. *r*, Pearson correlation coefficient; *P*, uncorrected two-tailed P value. No association reached nominal significance (all $P \geq 0.068$).

| Gene | Condition × Time | | | Cohen's d | CI Low | CI High |
|---|---|---|---|---|---|---|
| | *β* | *P* | FDR *q* | | | |
| CD14 | 0.185 | 0.163 | 0.852 | 0.288 | 0.034 | 0.580 |
| FCGR3A | -0.024 | 0.904 | 0.961 | -0.041 | -0.321 | 0.233 |
| CX3CR1 | -0.006 | 0.961 | 0.961 | -0.009 | -0.279 | 0.272 |
| CCR2 | 0.007 | 0.926 | 0.961 | 0.018 | -0.232 | 0.352 |
| SELL | 0.016 | 0.853 | 0.961 | 0.030 | -0.233 | 0.300 |
| CDKN1C | -0.039 | 0.853 | 0.961 | -0.048 | -0.304 | 0.231 |
| LST1 | 0.106 | 0.253 | 0.852 | 0.199 | -0.059 | 0.493 |
| HLA-DRA | 0.097 | 0.320 | 0.852 | 0.195 | -0.063 | 0.566 |

**Table S5. Condition-dependent expression changes in monocyte subset markers.** Canonical markers of classical (*CD14*, *CCR2*, *SELL*), intermediate (*HLA-DRA*), and non-classical (*FCGR3A*, *CX3CR1*, *CDKN1C*, *LST1*) monocytes were examined using LME models of $\log_2$(Transcripts per million; TPM) values (expression ~ condition × time + condition order + batch + sex + age + (1|subject); $n = 55$). $\beta$ and $P$ correspond to the condition × time interaction term (OT vs OC). FDR $q$ denotes BH correction across the eight markers. Cohen's $d$ represents the standardized within-participant difference in pre-to-post expression changes, with 95% confidence intervals estimated by participant-level bootstrap resampling (5,000 iterations). No marker showed a significant interaction (all FDR $q \geq 0.852$, all uncorrected $P \geq 0.163$), and all effect sizes were small. These findings provide no evidence of substantial condition-dependent changes in the expression of the selected monocyte subset markers.

| Pathway | mean ΔFC | *r* | *P* | |
|---|---|---|---|---|
| Nucleotide metabolism | -0.093 | -0.452 | 0.0005 | *** |
| Nucleotide import | -0.130 | 0.075 | 0.5873 | |
| Nucleotide synthesis and processing | -0.082 | -0.611 | 0.0000 | *** |
| Detoxification | -0.087 | -0.684 | 0.0000 | *** |
| ROS and glutathione metabolism | -0.084 | -0.505 | 0.0001 | *** |
| Organelle contact sites | -0.077 | -0.787 | 0.0000 | *** |
| Signaling | -0.070 | -0.373 | 0.0051 | ** |
| Calcium homeostasis | -0.080 | 0.112 | 0.4147 | |
| Mitochondrial permeability transition pore | -0.198 | -0.196 | 0.1509 | |

**Table S6. Correlations between genePC1 scores and differential expression of mitochondrial pathways showing significant condition effects.** The nine mitochondrial pathways with significant condition main effects (FDR $q < 0.05$; **Figure 2C**) are listed and grouped according to gene overlap, as shown in **Figure 2C**. For each participant, ΔFC was calculated as the difference in fold change in pathway (OT−OC) and mean ΔFC represents the average across participants. Pearson *r* represents the correlation between participants' pathway ΔFC and genePC1 scores (**Figure 4D**), and *P* denotes the two-sided nominal *P* value (**$P <$ 0.01, ***$P < 0.001$). Six of the nine pathways showed significant negative correlations with genePC1 ($P < 0.01$), all of which survived FDR correction across the nine pathways (FDR $q <$ 0.01).
